# Region-specific mechanosensation modulates *Drosophila* postural control behaviour

**DOI:** 10.1101/2025.04.22.645758

**Authors:** William Roseby, Jonathan A.C. Menzies, Victoria A. Lipscomb, Claudio R. Alonso

**Affiliations:** Department of Neuroscience, Sussex Neuroscience, School of Life Sciences, University of Sussex, Brighton BN1 9QG, UK

**Keywords:** Drosophila, larva, sensory neuron, self-righting, Hox genes

## Abstract

The relation between regional morphological features derived from the bilaterian body plan and the behaviours necessary to extract utility from such structures is not well understood. Here we use the *Drosophila* larva to investigate this ‘form-function’ problem focusing on the mapping of the regional stimuli that trigger an adaptive and evolutionarily conserved behaviour termed self-righting: a postural control system that allows the animal to restore its natural position if turned upside-down. Through the development of new methodologies that allow regionally-restricted mechanical stimulation and zonal-specific neuronal optogenetics, we find that multidendritic sensory neuron inhibition in anterior areas (thoracic/anterior abdominal) has a profound effect on self-righting performance, whilst inhibition of posterior sensory elements (mid and posterior abdomen) produces no effects. To gain insight into how regional neuronal inhibition affects the different subcomponents of the self-righting sequence we applied a deep neural network tracking method which revealed that reduction of neural activity in anterior sensory neurons primarily increases head casting behaviour, and that this, in turn, is strongly correlated with abnormally long self-righting times. Furthermore, to explore the mechanistic bases of our behavioural observations, we considered the hypothesis that the *Hox* genes – well known for their roles in axial developmental patterning – might play a role in the functional specification of multidendritic sensory neurons along the body axis. Molecular expression analysis of FACS-sorted neural populations, fluorescent immunolabelling and neuron-specific knock-down experiments demonstrate that normal sensory neuron expression of the *Hox* genes *Antennapedia* and *Abdominal-b* is necessary for self-righting in the *Drosophila* larva. Altogether, our work shows that region-specific mechanosensory processes mediated by multidendritic sensory neurons and instructed via *Hox* gene inputs are essential for self-righting, providing a link between regional structural features and an adaptive and widely evolutionarily conserved postural control behaviour.

## INTRODUCTION

Bilaterally symmetric animals (Bilaterians) represent the largest group of animals, including highly diverse phyla such as Arthropoda and Chordata. The bilaterian *bauplan* derives from the last common ancestor to all bilaterians, the urbilaterian [De Robertis & Sasai, 1996], which is believed to have possessed a substantial level of complexity featuring a nervous system, and a regionalised main body axis running from head-to-tail (antero-posterior, AP axis) as well as an orthogonal axis, the dorsoventral (DV) axis. Urbilaterians are thought to have lived in the ocean with a pelagic larval form and a benthic adult inhabiting the sea bottom, crawling around and burrowing in the rich nutritional environment provided by phytoplankton sediments [De Robertis & Tejeda-Muñoz, 2022]

Several functional features present in modern bilaterians are linked to the axial organisation derived from the urbilaterian AP and DV coordinates. For example, the anterior region in most bilaterians hosts arrays of sensory organs – eyes, olfactory and gustatory systems, and so on – all located in close proximity to the brain, presumably with the dual purpose of detecting new information as quickly as it is received when the animal moves forward and enters new environments, and to save ‘on wire’ and speed-up information transfer [Sterling and Laughlin, 2015]. Another salient feature is that the ventral side of the body is often specialised for contact with the substrate (or medium) through cilia, protruding appendages, limbs, or, simply equipped with morphological elements that provide some form of ‘grip’ or means to propel the animal forward as it moves around its habitat.

Nonetheless, for all these morphological specialisations to contribute to fitness, the animal must be capable of generating a suitable behavioural repertoire to safeguard an adequate body orientation in the face of unexpected environmental change. An example of this type of behaviour is a common postural control system termed *self-righting* [Picao-Osorio et al., 2015], which allows the animal to restore its normal orientation in respect to substrate should it find itself ‘upside-down’ [**Figure 1A**]. Self-righting is evolutionarily conserved all the way from insects to mammals [Ashe, 1970; Penn, 1995; Faisal & Matheson 2005; Jusufi et al., 2011; Feather-Schussler & Ferguson, 2016; Bagnall & Schoppik, 2018] [**Figure 1B**], including humans where it constitutes an element of medical tests evaluating motor development in young infants [McGraw, 1941; Teitelbaum et al., 1998; Siegel et al., 2024]. Notably, the relation between regional internal/external structural features derived from body plan organisation and the simple behaviours necessary to extract functionality from such elaborated structural features is not well understood. Here we study this ‘form-function’ problem in the *Drosophila* larva seeking to determine the signals that trigger self-righting and how such signals relate to body plan organisation and morphology.

**Figure 1:**
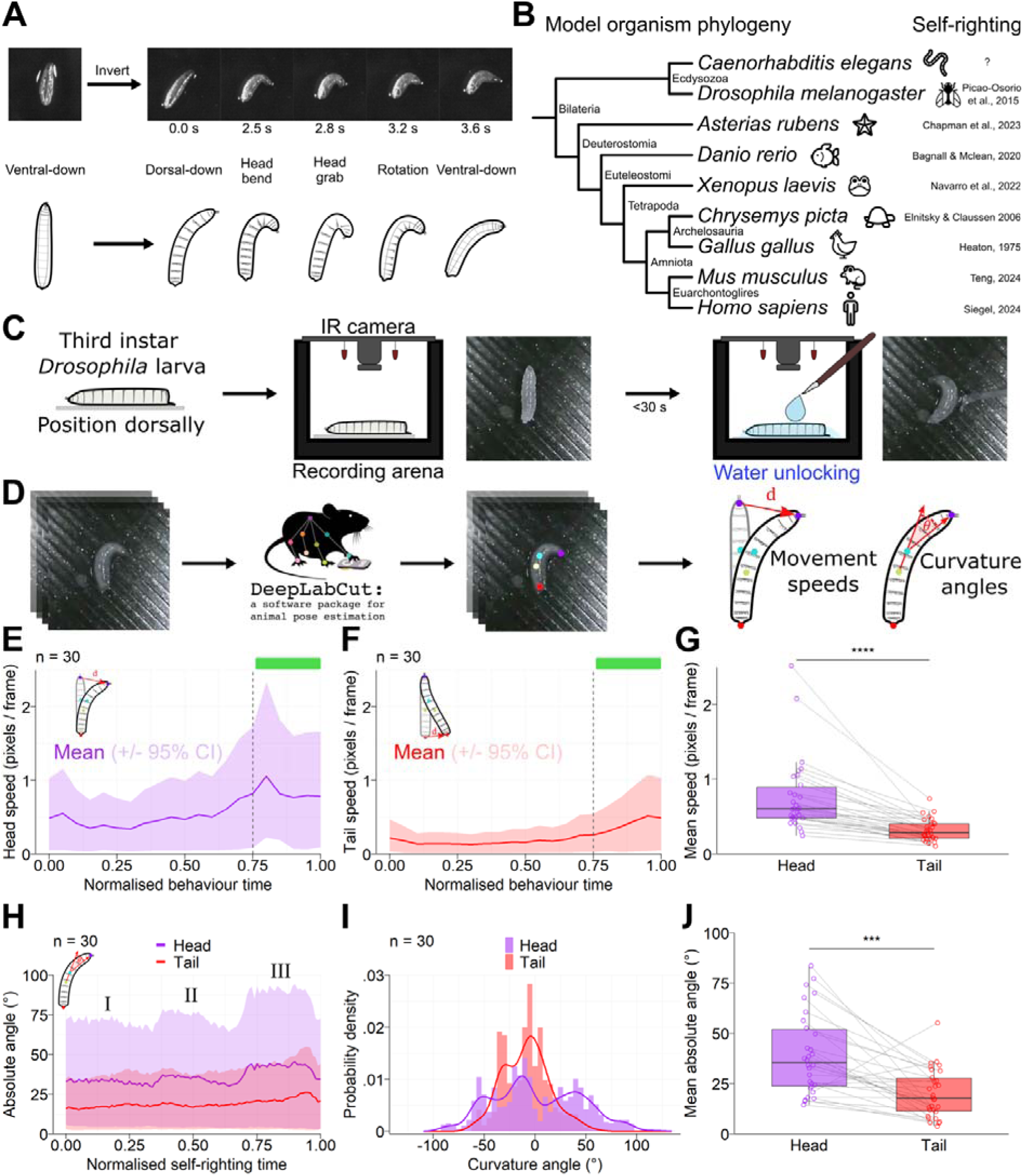
Quantitative analysis of self-righting behaviour. (**A**) Photographs (top) and diagrams (bottom) of the self-righting sequence of a first-instar *Drosophila* larva. After inversion of the normal posture, the self-righting sequence involves a 180° rotation of the body that begins at the head and takes around 3-5 seconds on average. **B)** A phylogenetic tree of common model organisms, with a reference to documented self-righting behaviour on the right. **C)** Experimental procedure for consistent recording of self-righting behaviour. Third-instar w^1118^ larvae were positioned on a dry coverslip with the dorsal side in contact with the surface before being placed in an arena for recording. Behaviour was ‘unlocked’ at the desired moment through the application of water with a moistened paintbrush. **D)** Extraction of behavioural features with video tracking. Video frames were analysed using DeepLabCut, where four points along the anterior-posterior axis were labelled. The coordinates of these points were used to calculate speed of movement as well as angles of body curvature. **E)** Mean head speed and **F)** tail speed over the course of recordings. The lines show the mean taken over all samples while the shaded areas indicate the 95% CI. The time course has been normalised to account for differences in the length of behaviour. The dotted line in E indicates where the self-righting sequence is predicted to begin on average, based on head movement. **G)** Mean speed of movement for the head and tail during the self-righting sequence. Points indicate mean speeds for individual samples, with measurements from the same larva being connected by lines. The box and whisker plots indicate the median, IQR and 1.5 IQR. ‘***’ = P < 0.001 for Wilcoxon signed rank tests, n = 30. **H)** Absolute angles of curvature of the head (purple) and tail (red) over the course of the self-righting sequence. The lines show the mean absolute angle while the shaded areas indicate the 95% CI. Due to the apparent curvature of the head in three discreet bursts, the head angles have been labelled I, II and III. **I)** The distribution of curvature angles for the head (purple) and tail (red), where negative values are left-handed bends and positive values are right-handed bends. The lines show a smoothed kernel density estimation for the probability density while the bars indicate counts binned to each 10°. **J)** Mean absolute angle for the head and tail during the self-righting sequence. Points indicate mean speeds for individual samples, with measurements from the same larva being connected by lines. The box and whisker plots indicate the median, IQR and 1.5 IQR. ‘***’ = P < 0.001 for Wilcoxon signed rank tests, n = 30. The median self-righting times before and after trimming based on head movement were 12.8 and 11.0 seconds respectively.

Previous work demonstrated that all motile developmental stages of the fruit fly display self-righting, including the three larval stages and the adult [Issa et al., 2021]; of these, the larval stage is probably the simplest to advance the question on stimulation given the lower morphological and neural complexity of the larva compared to adult forms. Further work identified several genetic elements essential for self-righting via activities in the sensory and motor domains [Picao-Osorio et al 2015; Picao-Osorio et al 2015; Issa et al. 2019; Klann et al. 2021]. Despite this progress, the signals that prompt the self-righting response remain largely unknown, except from the fact that gravity does not play a role [Klann et al., 2021].

To determine the stimuli that trigger larval self-righting we first developed a new approach that allows a precise temporal and quantitative analysis of self-righting responses, the ‘water unlocking technique’. Using this new technique in a series of combinatorial mechanical stimulation experiments providing specific surface contacts on the anterior, posterior, ventral and/or dorsal sides of the larva (and combinations therewith) we discover that for self-righting to occur the animal requires anterior dorsal contact in the absence of ventral anterior contact. Seeking to advance the understanding of the cellular processes that underlie the triggering of self-righting, we considered the possibility that sensory neurons might play a role; to test this, we conditionally inhibited different subpopulations of larval sensory neurons using a thermogenetic approach and determined that normal sensory neuron activity – in particular that of multidendritic sensory neurons – is required for normal larval self-righting to occur. Building on these observations, we developed another specialised approach – the ‘opto-axial method’ – which allows regional-specific optogenetic manipulation of sensory neurons located at different axial positions of the larva. Using the opto-axial approach we established that multidendritic sensory neuron inhibition in anterior areas (thoracic and anterior abdominal) has a profound effect on self-righting performance; in contrast, inhibition of posterior sensory elements (mid and posterior abdomen) has no noticeable impact on self-righting. Targeted application of this method shows the md subpopulation of daIVs also follows the axial pattern, albeit with reduced effect size compared to the entire md population. Exploring the genetic basis underlying these differential role of axial sensory neurons we explored the possibility that the Hox genes – known for their roles in axial developmental patterning [Mallo and Alonso, 2013] – might play a role; for this, we first demonstrated Hox expression in multidendritic sensory neurons, and, secondly, showed that normal sensory neuron expression of the Hox genes *Antennapedia* and *Abdominal-b* is necessary for “normal” self-righting in the Drosophila larva.

Altogether our work demonstrates that region-specific mechanosensory processes mediated by multidendritic sensory neurons and Hox gene inputs are essential for self-righting providing a link between regional internal structural features and an adaptive and widely evolutionarily conserved postural control behaviour. Based on the phylogenetic conservation of self-righting we propose that it may have been an ancestral behaviour present in the urbilaterian, suggesting that similar regional mechanosensory processes might play an important role in the triggering of self-righting in other animal systems.

## MATERIALS AND METHODS

### Fly stocks and maintenance

All fly stocks were reared on standard molasses food at 25 °C and 45% relative humidity, with a day-night cycle of 12:12. For experiments involving embryos or first instar larvae, stocks were transferred to a cage with an apple juice agar plate supplemented with yeast paste and samples were collected from the plate. For experiments with third instar larvae, samples were collected directly from the molasses food and briefly rinsed in water before being transferred to a 1% plain agar plate. To generate larvae for optogenetic experiments, crosses were carried out directly in molasses food supplemented with ATR. Most stocks were acquired from the Bloomington Drosophila Stock Centre (BDSC). For a full list of stocks used in this study, see Table 1 below.

**Table 1:** List of *Drosophila* stocks used in this study.

| Stock | Source | Reference |
| --- | --- | --- |
| *w<sup>1118</sup>* | Alonso lab | Hazelrigg et al., 1984 |
| *yw* | Alonso lab | Nash and Yarkin, 1974 |
| *yw; 109(2)80-Gal4* | BDSC #8769 | Gao et al., 1999 |
| *yw; 109(2)80-Gal4, UAS-mCD8::GFP* | BDSC #8768 | Gao et al., 1999 |
| *yw;; nompC-Gal4* | VDRC 2054 | Yan et al. 2013 |
| *w;; ppk-Gal4* | BDSC #32079 | Ainsley et al. 2003 |
| *w;; UAS-shi<sup>[ts]</sup>* | BDSC #44222 | Kitamoto, 2001 |
| *w; UAS-GtACR2* | BDSC #92984 | Govorunova et al., 2015 |
| *w;; 10XUAS-GCaMP6s::tdTomato* | Marta Zlatic | Grover et al., 2020 |
| *yv;; UAS-Antp- Valium10-RNAi* | BDSC #27675 | Perkins et al., 2015 |
| *yv;; UAS-Ubx- Valium10-RNAi* | BDSC #31913 | Perkins et al., 2015 |
| *yv;; UAS-AbdA- Valium10-RNAi* | BDSC #28739 | Perkins et al., 2015 |
| *yv;; UAS-AbdB- Valium10-RNAi* | BDSC #26746 | Perkins et al., 2015 |
| *yw;; UAS-Valium10-GFP* | BDSC #35786 | Perkins et al., 2015 |

### Behavioural assays

Unless otherwise stated, all behavioural experiments were conducted at an ambient temperature of 25.0 (+/− 1.0 °C) and a relative humidity between 40 and 60%. All experiments were conducted during a window of zeitgeber time ZT4-ZT8 (NB: ZT0 = lights turn on; ZT12 = lights off). Equal numbers of control and experimental larvae were tested under the same conditions on the same day, excluding occasional larvae that were discarded during the course of an experiment. Larvae were not distinguished by sex, so any sample was assumed to be a random mix of two sexes. For consistency between experimental setups, completion of self-righting behaviour was taken to be the time between experimental manipulation (e.g., water application), and the first moment when both tracheal trunks were visible along the whole length of the larva. Similarly, for completion of crawling, this was the time from experimental manipulation to completion of a peristaltic wave from posterior to anterior. Behavioural recordings were captured with a USB infrared CMOS camera (2MP OV2710, Arducam Technology Co. Ltd, Hong Kong) recording at 30 fps at 1920 × 1080. Behavioural arenas were 3D-printed in black PLA (1.75mm, RS Pro, UK) using a Prusa MK3S (Prusa Research, Czech Republic), with a layer height of 0.2 mm and 15% infill.

#### Water unlocking technique

For several experiments, we required a method of controlling larval position and restricting movement prior to beginning a behavioural test. We developed a technique centred around the presence of moisture, which is required for larval movement. We restricted movement by first drying larvae on a soft tissue, then subsequently transferring them to a standard microscope coverslip (22 × 22 mm, 0.13 − 0.17 mm thickness; Thermo Fisher Scientific, USA) using a paintbrush. The desired position (e.g., dorsal side down) was achieved by gently rolling the larva with a paintbrush.

We restored the capacity for normal movement through the delivery of water with a moistened paintbrush that had most of the bristles removed. The paintbrush was brought to the interface of the coverslip and the larval midpoint, which allowed larval hydration while preventing unwanted movement due to water adhesion and surface tension. The time from larval drying to water delivery was kept to a minimum (in all cases < 30 s) to avoid larval desiccation. This technique was often used as an alternative to the standard self-righting assay which involves rolling a larva directly on agar, and we refer to it in other sections as the ‘water unlocking technique’.

#### Contact based assays

The effect of localised substrate contact was tested by placing foraging third instar w^1118^ larvae onto a substrate with specific portions of the body touching the surface. In the dorsal contact experiments, we positioned larvae on a coverslip such that only the dorsal side contacted the surface. This involved one of three positions: the anterior half, posterior half, or whole dorsal side touching the surface. These conditions were chosen in a randomised order for each larva. After positioning, larvae were released using the water unlocking technique and we measured the occurrence and timing of self-righting within the subsequent 60 s.

For the ventral contact experiments, larvae were placed ventral side down on a coverslip, with the anterior half, posterior half or whole ventral side in contact with the surface. The larva on the coverslip was then placed dorsal side down onto a 1% agar plate, such that the whole dorsal side contacted the agar. This provided sufficient moisture to facilitate movement. We used a 3D-printed coverslip holder of 0.5 mm height to provide consistent contact to the dorsal and ventral sides of the larva. After placement, we noted the occurrence of the first behaviour. If the larva performed enough peristaltic waves to elicit a translation of more than one segment, the behaviour was labelled as crawling, even if the larva later performed self-righting.

#### Thermogenetic inhibition of sensory neurons

To conditionally inhibit sensory neuron populations, we used *shibire^[ts]^ (shi^[ts]^)*, a temperature-sensitive allele of the shibire protein which is required for synaptic vesicle recycling. Exposure of *shibire^[ts]^* to restrictive temperatures above 30 °C results in synaptic vesicle depletion and cessation of synaptic transmission. We generated *shibire^[ts]^* heterozygote larvae by crossing male flies carrying *UAS-shi^[ts]^*with virgin females carrying a sensory domain Gal4. We collected six first instar larvae at a time and allowed them to acclimate on 1% agar at 25 °C for 5 min. We tested self-righting in the typical fashion by rolling each larva onto its dorsal side with a small paintbrush and measuring the time to the restoration of typical posture, up to a maximum of 300 s. We then transferred the agar onto a Peltier device (TC-36-25, TE Technology Inc, USA) pre-heated to 32 °C and left the larvae for 20-30 min for *shibire^[ts]^* inhibition to occur. We tested self-righting again as above, before removing the agar from the Peltier, allowing a recovery of 45 – 60 min, and testing self-righting a final time. Prior to each self-righting test, larvae were gently rolled a few times, with the aim of depleting any remaining synaptic vesicles during the *shibire* inhibition. We tested control larvae homozygous for *UAS-shi^[ts]^* in an identical fashion. While we started with equal sample sizes for controls, some larvae were discarded during the experiment due to wandering onto the Peltier device, leading to some small imbalances in sample sizes between the groups.

#### Localised optogenetic inhibition of sensory neurons (Opto-axial approach)

To locally inhibit groups of sensory neurons, we developed an optogenetic approach using physical masking of light to sets of segments along the anterior-posterior axis. We generated heterozygote larvae expressing GtACR2 in multidendritic sensory neurons by crossing male flies carrying *UAS-GtACR2* with virgin females carrying *109(2)80-Gal4* or *ppk-Gal4*. Crosses were carried out in food supplemented with ATR to a concentration of 5 mM. We selected foraging third instar larvae and allowed them to acclimatise on 1% agar for 5 min. To test a larva, we placed it dorsal side down on a coverslip and placed it in a custom 3D-printed arena. The arena contained a slit in the base measuring 8 x 1 mm through which light was delivered. We used a 1 W LED with a peak wavelength of 470 nm (OSCONIQ P3030, Intelligent LED Solutions, UK) to inhibit neurons expressing GtACR2. The LED was attached to the underside of the arena and delivered ∼2.12 mWmm^−2^ to three segments of the larva positioned above. The LED was activated manually after positioning the larva via a switch. After light delivery, the larva was quickly released using the water unlocking technique. The subsequent behaviour was recorded up until the completion of self-righting. Larvae were subjected to five illumination conditions in a random order, with light targeting four sets of three segments from anterior to posterior (T1-T3, A1-A3, A4-A6, A7-A9), and one control condition where the light was not activated (None). Control genotypes were generated by crossing one parental line with a line carrying the background genetics of the other line (i.e., w or yw), and were tested in an identical fashion. The expression strengths of the 109(2)80-Gal4 and ppk-Gal4 lines were compared via crossing these to the line *w; +; 10XUAS-GCaMP6s::tdTomato*, then measuring fluorescence intensity of tdTomato in third instar progeny. The tdTomato fluorescence intensity was quantified as the mean grey value of an equal-sized ROI drawn around the daIV neuron ddaC, using the red channel of images acquired on a Leica DM6000. All larvae were imaged from the dorsal side. The mean fluorescence intensity was almost equal, suggesting equal expression strength within ddaC across the two drivers. The GCaMP signal was not measured.

### Behavioural feature quantification

#### DeepLabCut implementation

To quantify features of self-righting behaviour, we first carried out tracking of larval body positions using the pose estimation machine learning tool DeepLabCut [version 2.2.3, Mathis et al., 2018]. Using recordings with a top-down view, we labelled four points along the anterior-posterior axis. This included the tip of the head, the tip of the tail, and two points close to the midpoint of the larva separated by one segment. Points were always labelled along the apparent midline of the larva, irrespective of its rotation with respect to the camera. For the training dataset, we labelled 100 frames extracted using k-means. Training was carried out with default settings (ResNet50) for 500,000 iterations. Analysis and video generation were also performed with default settings. Network performance was assessed by visual examination of videos, which showed accurate labelling in all cases. All videos input to DeepLabCut were manually trimmed so that the start and end coincided with the start and end of the behavioural sequence. All training and analyses were conducted on a PC running Windows 10 Home and using an NVIDIA RTX 2070 Super (NVIDIA, USA).

#### Feature calculation

Body coordinates retrieved using DeepLabCut were used to calculate behavioural features. All data wrangling and analysis was carried out in R [version 4.2.2., R Core Team, 2024]. Speed was calculated as the two-dimensional Euclidean distance travelled over each frame. Curvature angles were calculated using trigonometric functions on the vectors between the larval midpoints and the end points. Here, we extended the vector between the two midpoints to acquire a ‘straight’ vector. We used this as an approximation of where the head (tail) position would be if the larva had no curvature. In combination with vectors describing the position of the head (tail) with respect to the midpoint, curvature was calculated as:

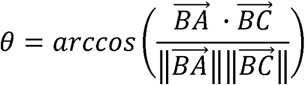

Where BA is the vector from midpoint to head (tail) and BC is the vector from midpoint to the ‘straight’ head (tail). To calculate if curvature left- or right-handed, we first normalised all coordinates by rotating them with respect to the video frame, with the degree of rotation applied to align the two midpoints along the video y-axis. Left and right positions were then designated on the basis of x-axis values of the head (tail) with respect to the midpoint, with lower x values representing right-handed curvature and greater x values representing left-handed curvature. Changes in curvature direction where then calculated as the number of times the x-axis value of the head (tail) went from increasing to decreasing or vice versa.

### Immunohistochemistry

Immunohistochemistry of 109(2)80>GFP embryos was carried out as previously described [Klann et al., 2021]. Embryos were collected at around 15 hours after egg laying, corresponding to developmental stage 16, to ensure sufficient development of the PNS. Dechorionation was carried out in 100% bleach for 2 min and samples were fixed for 30 min in a solution of 10% formaldehyde and heptane at a 1:1 ratio. Following an overnight incubation in methanol, embryos were rehydrated with 50% v/v methanol and PBS supplemented with 0.3% v/v Triton-X (henceforth PTBX). Embryos were then incubated with the primary antibody solution overnight at 4 °C. After washing in PBTx, the secondary antibody incubation was carried out for 2 hours at room temperature. For concentrations of primary and secondary antibodies, see Table 2.

**Table 2:** Antibodies used in immunohistochemistry. All solutions were prepared in PBS supplemented with 0.3% v/v Triton-X.

| **Antibody** | **Type** | **Host animal** | **Source** | **Dilution** |
| --- | --- | --- | --- | --- |
| anti-GFP<br>(ab13970) | Primary | Chicken | Abcam | 1:100 |
| 22C10 (anti-Futsch) | Primary | Mouse | DHSB | 1:10 |
| anti-Antp 4C3 | Primary | Mouse | DHSB | 1:100 |
| anti-Abd-B<br>1A2E9 | Primary | Mouse | DHSB | 1:100 |
| Alexa-Fluor anti-chicken A488 | Secondary | Goat | Invitrogen | 1:500 |
| Alexa-Fluor anti-mouse A555 | Secondary | Donkey | Invitrogen | 1:500 |

For imaging, embryos were then mounted on a microscope slide in a solution of 75% v/v glycerol in PBS. Imaging was carried out on a Leica SP8 scanning confocal microscope. Laser power, scanning routine, aperture size, sensor gain and offset, and z-slice step size were varied depending on the sample to achieve optimal brightness, contrast and resolution in each channel. Image manipulation was carried out in FIJI, which involved image rotation, further adjustment of channel brightness and contrast, and z-projection generation.

### Cell sorting and RT-PCR

Collection of multidendritic neurons from 109(2)80>GFP larvae was performed as previously described [Klann et al., 2021], following a protocol described by Harzer and colleagues for the collection of neuroblasts [Harzer et al., 2013]. Briefly, this involved the opening of newly hatched first instar larvae at different points along the body axis, before tissue dissociation using collagenase I and papain. Following washing with Rinaldini’s solution and Schneider’s medium, the cells were homogenised and taken for fluorescence-activated cell sorting (FACS) on a BD FacsMelody (BD Biosciences). GFP-expressing cells were collected directly into TRIzol and homogenised via vortexing, and non-fluorescent cells were discarded. Around 50 larvae were processed per replicate resulting in the collection of around 5,000 cells.

Cells collected via FACS were immediately used for RNA extraction following a standard phenol-chloroform extraction from TRIzol. Phase-separated RNA was precipitated in isopropanol and glycogen before being resuspended in RNase-free water. RNA was reverse transcribed to cDNA using the SuperScript IV kit (Invitrogen) using oligo dT(20) primers. cDNA was then amplified via polymerase chain reaction (PCR) using standard Taq DNA polymerase (NEB) for 40 cycles and with annealing temperature of 60 °C. Primers were included at a final concentration of 0.4 μM for the Hox genes antennapedia, ultrabithorax, abdominal-A and abdominal-B, as well as for the housekeeping gene rp49, the sequences of which can be found in Table 2. Following amplification, gel electrophoresis was carried out on a 2% agarose gel containing ethidium bromide and submerged in TAE buffer. The amplified DNA was run alongside a 100 bp ladder (NEB). Control samples were also run which either did not undergo reverse transcription (no RT), or lacked the cDNA in the PCR step (no template), to account for genomic DNA or non-specific amplification respectively.

**Table 3:**
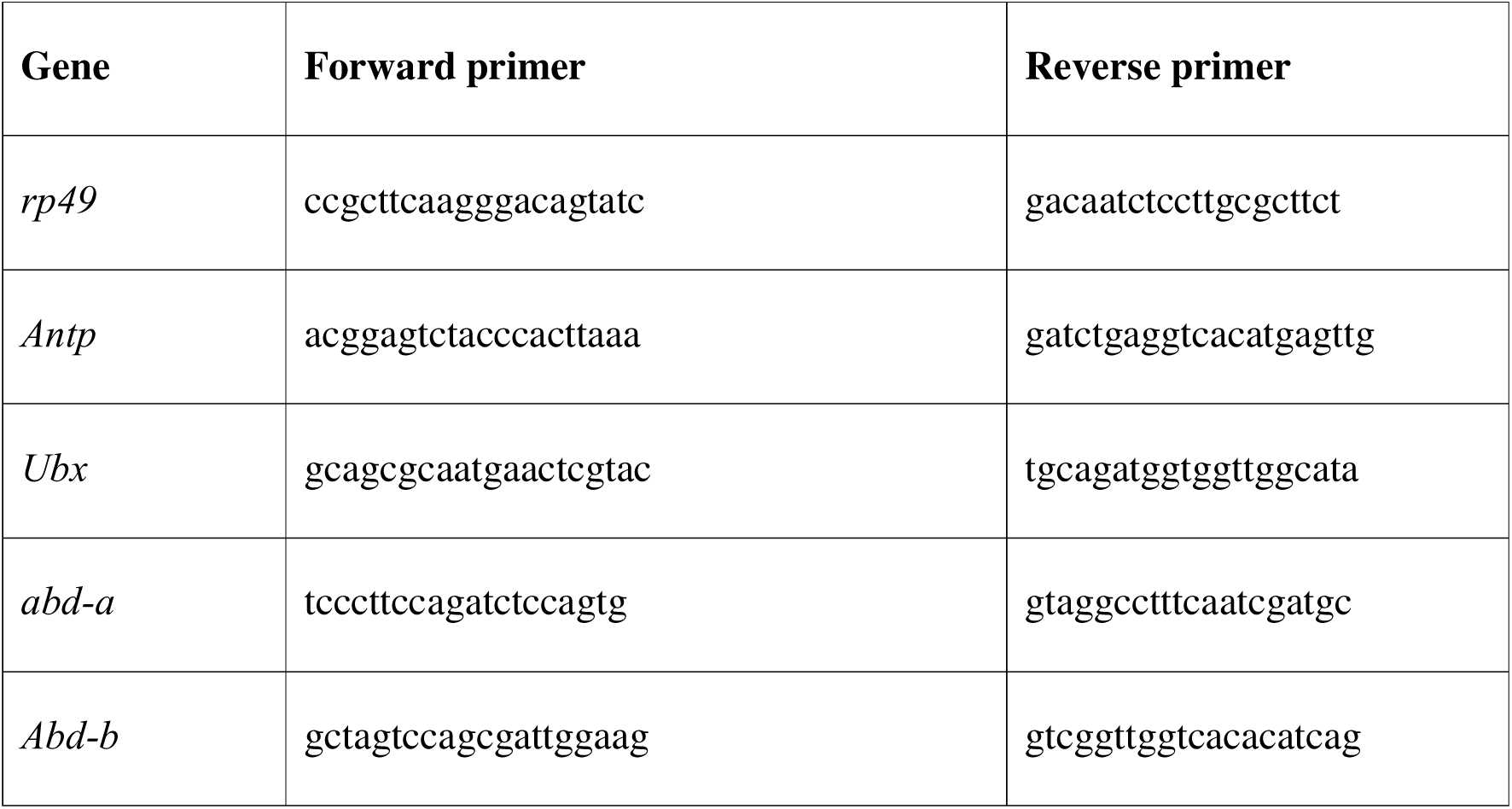
Sequences of primers used in RT-PCR.

### Statistical analysis

All statistical analysis was carried out in R (version 4.2.2). Continuous response data were first checked for normality through visual examination of distributions as well as with the Shapiro-Wilk test. In the case of non-normal data, paired comparisons between experimental conditions were conducted using the Wilcoxon signed rank test. Relationships between continuous data (and count data) were assessed with Spearman correlations.

For continuous data predicted by multiple factors, such as self-righting times under localised optogenetic inhibition, we fit linear mixed effects models. Data were log-transformed to improve model fit. If a given predictor had a single control level (illumination condition), we used treatment contrasts; otherwise, effects contrasts were used (genotype). Fixed effects were estimated for experimental manipulations (genotype, illumination condition) while random intercepts were estimated to account for sample effects. Model fit was assessed by visual examination of residual and random effect distributions as well as numerical examination with the Shapiro-Wilk test; model fit was ideal in all cases. P-values for marginal model effects were calculated following analysis of deviance with F-tests. In the case of a significant interaction effect, we conducted post-hoc contrasts of estimated marginal group means, conditional on genotype. To reduce the number of post-hoc tests, we followed Dunnet’s approach to compare each experimental condition with the control condition. One-sided tests were used in the case of a specific hypothesis of response increase or decrease. P-values were adjusted with Sidak’s correction for multiple comparisons.

Count data predicted by multiple factors, such as the number of head casts during self-righting, were modelled in a similar fashion as above but using negative binomial models. Model fit was assessed by calculating the ratio of residual deviance to degrees of freedom of the residuals. P-values for marginal model effects were calculated following analysis of deviance with X^2^ tests. Post-hoc comparisons were carried out as described above for continuous data.

Matched binary data, such as the occurrence of self-righting under localised substrate contact, were tested with Cochran’s Q test. The test was first applied across conditions to infer the existence of group differences, before being applied pairwise between groups (where it is equivalent to the McNemar test) to determine the location of the significant differences.

## RESULTS

### Self-righting behaviour involves greater speed and curvature of the head than tail

A simple visual characterisation of self-righting behaviour (as has been done previously, see Picao-Osorio et al., 2015) suggests that the sequence involves differential movement of body regions along the anterior-posterior axis over time [**Figure 1A**], an observation that has been supported quantitatively by a study employing physical modelling of larval movement [Loveless et al., 2021]. Furthermore, observation of this behaviour in a variety of model animals – across which it is highly conserved, including in humans [**Figure 1B**] – also points to a particular involvement of the anterior region of the body in initiating the first stages of rotation that is key to the proper execution of the sequence. However, it remained unclear how these regional differences might relate to the specific sensory conditions that induce self-righting. To ratify previous findings, and establish a framework for further behavioural analyses, we sought to develop a quantitative account of typical self-righting behaviour. To this end, we established a novel method for controlling larval posture we call the ‘water-unlocking technique’, which enabled us to record self-righting behaviour in a controlled and reproducible manner. This method involved the positioning of a third-instar w^1118^ larva in the dorsal-down posture on a dry glass surface. The induction of self-righting was then elicited by the delivery of a water droplet, which permitted the larva to move freely while behaviour was recorded [**Figure 1C**]. To generate data on the posture of larvae throughout self-righting, we analysed recordings with DeepLabCut, a deep learning algorithm to track body coordinates over time [Mathis et al., 2018]. We labelled four points along the AP axis – head, tail and two middle points – from which we calculated speed and curvature angle of the head and tail [**Figure 1D**].

When examining head and tail speed over the course of entire recordings, we noticed a similar trend for both body parts, involving an initial lull of activity that increased towards the end of the recording [**Figure 1E, F**]. Since we wanted to focus on the self-righting sequence itself, we subset the data based on head movement to exclude the initial lull phase [green bar, **Figure 1E**]. As expected, we observed that speed during the self-righting sequence was significantly greater for the head than for the tail [**Figure 1G**], confirming the notion that the anterior plays an important role. Since lateral bending of the body is also a key part of the self-righting sequence [Picao-Osorio et al., 2015] we also examined the trends of absolute curvature of the head and tail over time. Interestingly, the average curvature for the head appeared to follow three distinct phases, characterised by a relatively steady degree of curvature that quite suddenly increased to a higher degree of steady curvature [**Figure 1H**, purple]. We suggest this observation likely reflects a repeated back-and-forth bending of the head that occurs prior to the rotational aspect of self-righting, which only occurs after opportunities for surface attachment have been exhausted. In contrast, the tail showed no such changes in curvature, with the curvature angle only subtly increasing in magnitude over the course of the behaviour [**Figure 1H**, red]. To better understand the distribution of curvature angles, as well as their directionality, we examined the probability densities for relative head and tail angles. For the tail, angles were somewhat normally distributed around zero, albeit with a small hump at 30° to the left [**Figure 1I**, red], suggesting that most tail angles are quite small, although there may be a slight preference for left-handed bends. Opposingly, the distribution for head angles was centred slightly left of zero, and showed distinct peaks at around 50° to both the left and the right [**Figure 1I**, purple]. This indicates that while head curvature is stronger than that of the tail, it tends to more often occupy a specific range of moderate curvature at around 50°. While these angle distributions suggested a possible chirality in the curvature occurring during self-righting, which showed a slight group-level preference for left-handed bends [Supplementary material **Figure S1**]. Direct comparison of the absolute curvature angles confirmed that head angles were indeed significantly greater than tail angles [**Figure 1J**], indicating that self-righting involves a particularly strong bending of the head. Overall, these results indicate that unlike other behaviours such as rolling [Cooney et al., 2023], self-righting is characterised primarily by movement and bending of the head, which tends to follow a profile of increased but moderate bending over time.

### Larval self-righting requires anterior dorsal contact but is abolished by ventral anterior contact

Our initial analysis of self-righting behaviour revealed a clear preference for anterior movement and curvature over that of the posterior. Because different behavioural sequences are often related to the presence of specific sensory stimuli, we questioned to what extent this asymmetry in muscle contractions was related to spatial differences in sensory processing. While it was apparent that self-righting depends on substrate contact at the dorsal side of the body, and not gravity [Klann et al., 2021], it was unknown if sensory stimuli at different areas along the AP axis would result in different behavioural outcomes. To tackle this problem, we devised a set of experiments involving the presence of substrate contact at restricted regions along the AP axis. We again used the water-unlocking technique in order to precisely position larvae in the desired posture [**Figure 2A**]. We placed each third instar w^1118^ larva in three positions with dorsal substrate contact: whole body contact, the anterior half only, or the posterior half only [**Figure 2B**], and observed the occurrence of self-righting. As expected, whole body contact led to the performance of self-righting as usual. Interestingly, while the majority of larvae also performed self-righting in the anterior contact position, the posterior contact condition failed to elicit self-righting in most larvae [**Figure 2C**], suggesting anterior substrate contact is required for self-righting. While we observed no clear visual differences self-righting sequence between the whole body and anterior only conditions, comparison of self-righting times suggested a slight trend towards faster times in the anterior-only condition [**Figure 2D**]. We suspect this is due to reduced friction experienced by larvae in the anterior contact condition, allowing them to complete the behaviour slightly faster than with whole body substrate contact.

**Figure 2:**
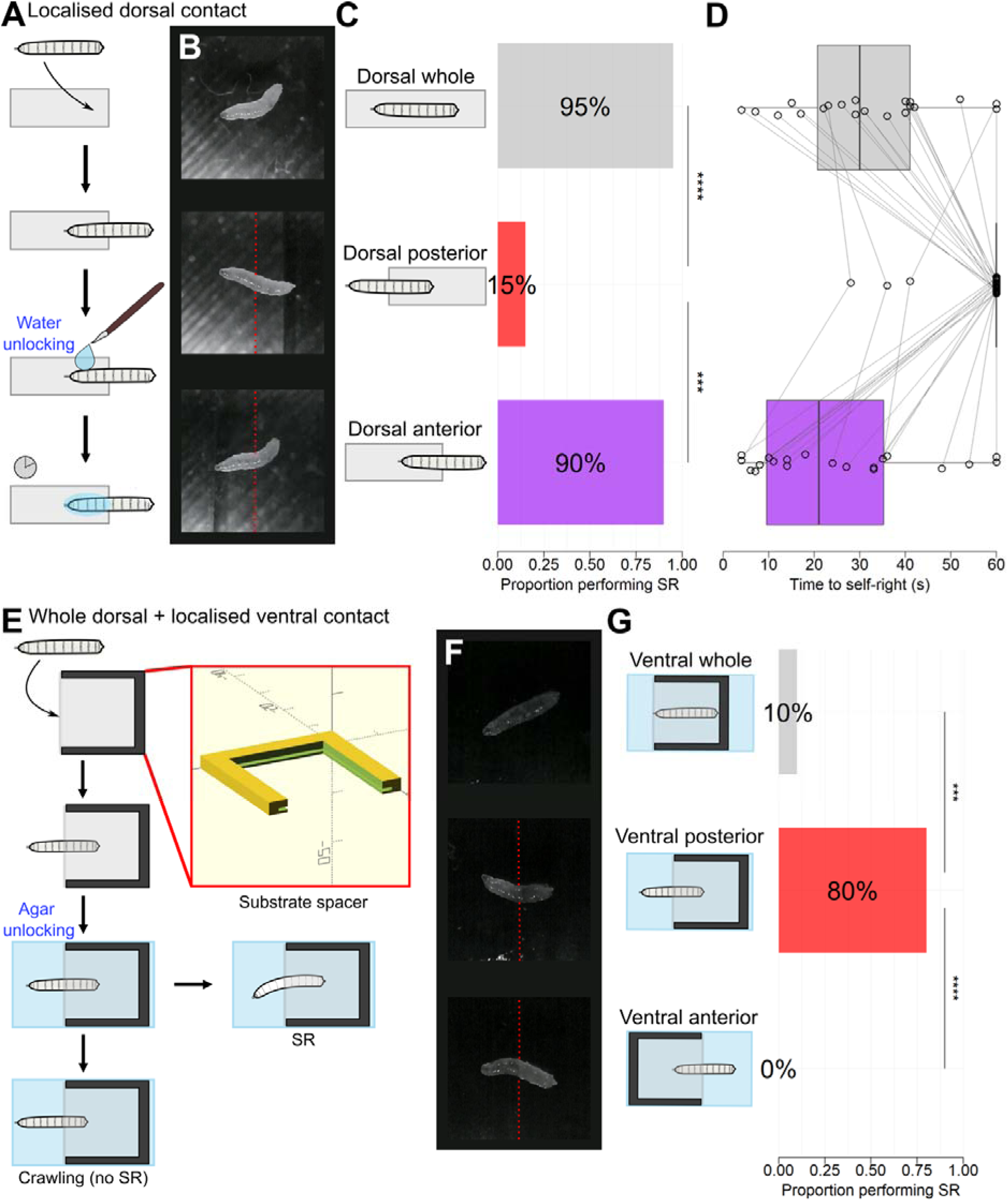
Effects of localised substrate contact on self-righting. **A)** Experimental procedure for investigating localised dorsal substrate contact. Third instar w^1118^ larvae were first placed dorsal side down on a dry glass coverslip in the desired position. Movement was then unlocked via the application of water with a paintbrush. **B)** Photographs of larvae in three conditions of dorsal substrate contact. The red dashed line indicates the boundary of the coverslip. **C)** Proportion of larvae that performed self-righting in the three conditions of dorsal substrate contact. Group comparison: Cochran’s Q_(2)_ = 28.35, P < .001, n = 20. ‘***’ = P < .001, ‘**’ = P < .01 for pairwise Cochran Q tests. **D)** Self-righting times in the three conditions of dorsal substrate contact. For larvae that didn’t self-right within 60s, the time is shown as 60. Points show times for individual tests, with measurements from the same larva being connected by lines. The box and whisker plots indicate the median, IQR and 1.5 IQR. P = 0.047 for a Wilcoxon signed rank test comparing the anterior and whole-body conditions, n = 17. **E)** Experimental procedure for investigating the combination of whole-body dorsal substrate contact with localised ventral substrate contact. A coverslip was held inside a custom 3D-printed mount (red box) to provide consistent contact. Third instar larvae were placed ventral side down on the coverslip in the desired position. The coverslip and larva were then placed onto agar, providing whole-body dorsal contact and sufficient moisture for movement. Larvae generally performed crawling or self-righting. **F)** Snapshots of larvae in the three conditions of ventral substrate contact. The red dashed line indicates the boundary of the coverslip. **G)** Proportion of larvae that performed self-righting in the three conditions of ventral substrate contact. Group comparison: Cochran’s Q_(2)_ = 29.50, P < .001, n = 20. ‘****’ = P < .0001, ‘***’ = P < .001 for pairwise Cochran Q tests. The mean self-righting time for larvae that performed self-righting with ventral contact was 32.8 seconds.

Given the apparent role of localised dorsal contact in eliciting self-righting, we then questioned if self-righting can still occur under a combination of dorsal and ventral substrate contact. Specifically, we were primarily interested to test if any differences would be observed with ventral contact present at different regions along the AP axis. To test this, we arranged larvae ventral side down on the substrate. We then subsequently placed the larvae dorsal side down onto an agar substrate, providing whole body dorsal contact and permitting movement from the moisture of the agar. We used a 3D-printed spacer to provide consistent contact to the two sides of the larva [**Figure 2E**]. As with the dorsal contact experiments, we placed larvae in three positions of ventral contact: whole body, anterior half, or posterior half only [**Figure 2F**]. Across these conditions, we observed an inverse effect to that of the localised dorsal contact: while the posterior only ventral contact permitted self-righting in most cases, both whole body and anterior-only ventral contact largely prevented the occurrence of the behaviour [**Figure 2G**]. These latter conditions instead elicited crawling in the majority of larvae, suggesting that anterior ventral contact is sufficient to suppress self-righting in favour of crawling despite the continued presence of dorsal substrate contact. In combination, these results highlight the anterior of the larva as a key region for the modulation of behaviour through mechanosensory stimulation.

### Conditional inhibition of multidendritic neurons delays self-righting

Although our results thus far suggested that self-righting depends on mechanical stimulation at the anterior, it remained unknown how this stimulation was received by mechanosensory neurons. Therefore, we decided to explore how self-righting behaviour correlated with the activity of localised mechanosensory neurons. However, it also remained unclear what populations of sensory neuron types were responsible for detecting substrate contact. While previous research suggested a role of the multidendritic neurons – in particular the highly branched daIV neurons – this relied on constitutive inhibition of these cells throughout development [Klann et al., 2021], which could have produced off-target knock-on effects on the rest of the nervous system [Fushiki et al., 2013; Kaneko et al., 2017]. Thus, we wished to implement a conditional approach to temporarily inhibit various groups of sensory neurons, which display a stereotyped arrangement along the AP axis [**Figure 3A**]. To achieve this, we used the Gal4/UAS system to express a temperature-sensitive allele of *shibire* (*shi^[ts]^*) in subpopulations of sensory neurons, which impairs synaptic transmission when exposed to temperatures greater than 30 °C [Kitamoto, 2001; Kasuya et al., 2009]. We tested self-righting behaviour under synaptic inhibition by measuring the time taken by first instar larvae to right themselves on an agar substrate after being rolled over with a paintbrush. To control for the effects of *shi^[ts]^* at the restrictive temperature of 32 °C, we also assayed self-righting before and after this inhibition at a permissive temperature of 25 °C [**Figure 3B**].

**Figure 3:**
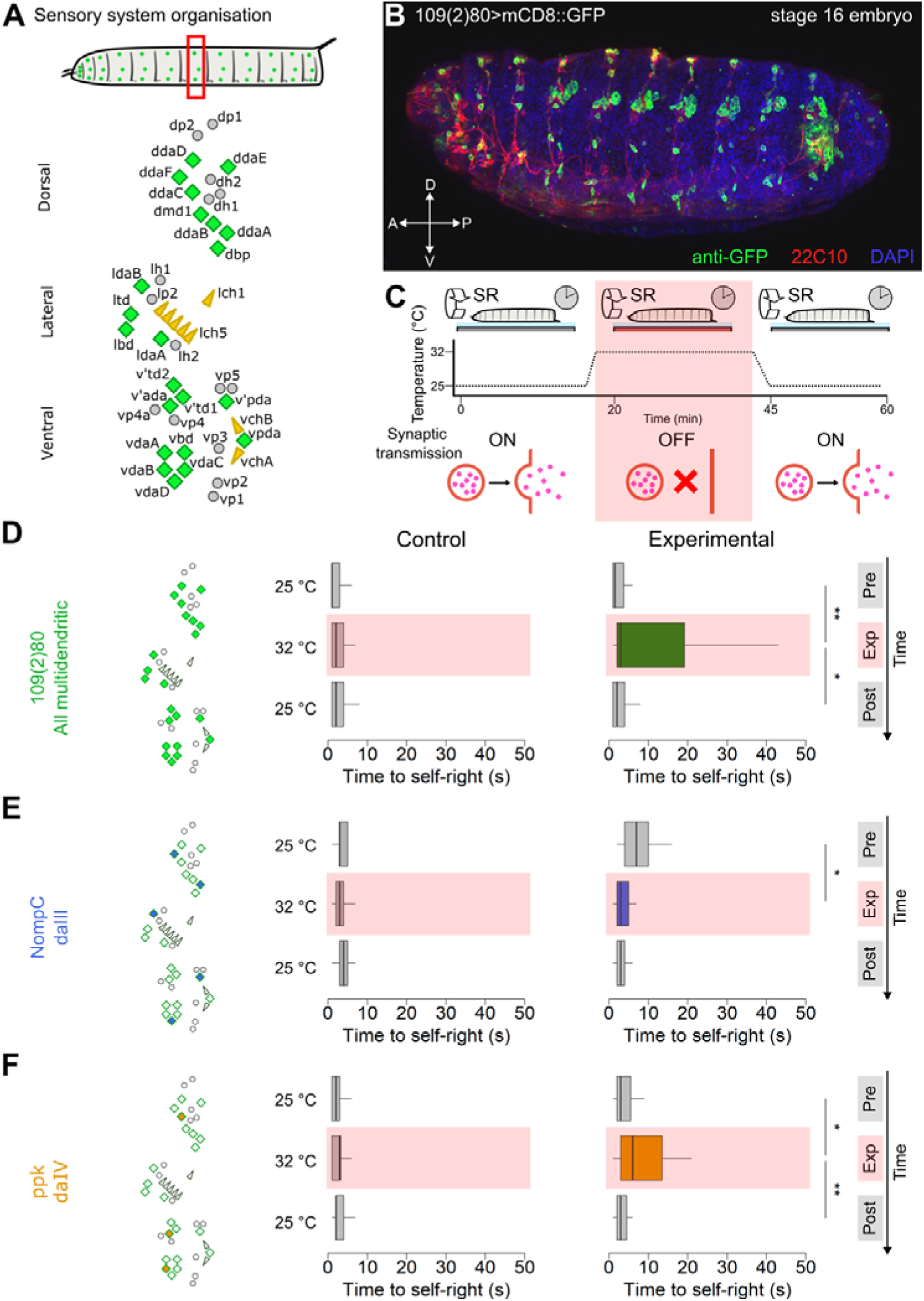
Conditional thermogenetic inhibition of multidendritic sensory neurons. **A)** The peripheral nervous system of the *Drosophila* larva. The diagram (top) shows the larva with the segmentally repeated clusters of sensory organs. The canonical hemisegmental arrangement (bottom) shows the es organs (grey circles), chordotonal organs (yellow triangles) and multidendritic neurons (green diamonds). Adapted from Orgogozo & Grueber (2005). **B)** A confocal z projection of a stage 16 embryo with GFP expression in the 109(2)80-Gal4 domain. The embryo has been immunolabelled with anti-GFP (green), 22C10 (axonal tracts, red) and DAPI (nuclei, blue). **C)** Experimental procedure for conditional inhibition of sensory neurons expressing *shibire*^[ts]^ (*shi^[ts]^*). Self-righting is first performed with first instar larvae at 25 °C, at which shibire is functional and synaptic transmission occurs normally. The substrate is then heated to 32 °C and self-righting is tested again. This temperature impairs shibire function and inhibits synaptic vesicle recycling and release. The temperature is then lowered back to 25 °C allowing restoration of shibire function. Self-righting is tested again to ensure recovery from the conditional inhibition. **D-F)** Self-righting times for first-instar larvae expressing shibire in different sets of sensory neurons. In each case, the location of the sensory neuron population within the hemisegmental arrangement is shown (left). Box and whisker plots indicate self-righting times in each temperature condition, for *UAS-shi^[ts]^* controls (left) and age-matched experimental genotypes (right). The temporal order of the temperature conditions follows from top to bottom of each plot. ‘*’ = P < .05, ‘**’ = P < .01, ‘***’ = P < .001 for pairwise Wilcoxon signed-rank tests. Individual observations are not displayed for visualisation purposes due to large variance. **C)** Self-righting times of *109(2)80>shi^[ts]^* larvae, expressing *shi^[ts]^* in all multidendritic neurons. n_[exp]_ = 38, n_[control]_ = 37. **D)** Self-righting times of *NompC>shi^[ts]^* larvae, expressing *shi^[ts]^* in daIII neurons. n_[exp]_ = 17, n_[control]_ = 17. **E)** Self-righting times of *ppk>UAS-shi^[ts]^* larvae, expressing *shi^[ts]^* in daIV neurons. n_[exp]_ = 39, n_[control]_ = 40.

Given the previous finding of a contribution of multidendritic (md) neurons to self-righting, we first tested inhibition in the 109(2)80-Gal4 domain, which drives expression in all multidendritic neurons [Gao et al., 1999] as well as in some oenocytes and chordotonal organs [Grueber et al., 2002], and also a few cells in the CNS [Hughes & Thomas, 2007]. Because md neurons in general comprise several classes of sensory neurons, their testing emerged as a suitable starting point for our conditional inhibition analysis.

Our results show a significant increase in self-righting times at the restrictive temperature, confirming the suggestion that multidendritic neurons play a role in self-righting. However, the entire set of multidendritic neurons contains several classes of neurons that detect both external mechanical stimuli (daII, daIII and daIV) [Hwang et al., 2007; Tsubochi et al., 2012; Yan et al., 2013] as well as proprioceptive feedback (daI, bd) [Hughes & Thomas, 2003; Vaadia et al., 2019]. To narrow down the multidendritic population, we also tested other Gal4 lines with class-specific expression domains. We initially reasoned that since substrate contact is an innocuous mechanical stimulus, it might depend on the daIII neurons, which are known to respond to gentle touch [Yan et al., 2013]. Thus, we tested the conditional inhibition in the NompC-Gal4 domain, which drives expression in the daIII neurons. Contrary to our hypothesis, self-righting was not delayed at the restrictive temperature; in fact, there was a trend towards faster self-righting times at this temperature [**Figure 3D**], suggesting that daIII neurons are not vital for detecting the substrate contact that induces self-righting. An alternative population of sensory neurons is the highly branched daIV neurons. This population have been strongly implicated in nociception due to their involvement in rolling behaviour [Tracey et al., 2003; Hwang et al., 2007; Ohyama et al., 2013]; however, they have also been linked to sensory feedback during crawling behaviour [Ainsley et al., 2003; Gorczyca et al., 2014; Jang et al., 2019], as well as to self-righting [Klann et al., 2021]. In agreement with previous findings, we indeed found that inhibition of daIV neurons via the ppk-Gal4 domain caused substantial delays in self-righting times [**Figure 3E**], suggesting that these neurons are indeed important for normal self-righting behaviour. However, the effect size was smaller than that for the entire multidendritic population, suggesting neurons other than the daIVs are important for self-righting. Since they have been implicated in other behaviours like rolling and crawling [Caldwell et al., 2003; Ohyama et al., 2015], we also examined the effect of chordotonal organ (CO) inhibition on self-righting. While we did observe a strong effect on self-righting times [supplementary material **Figure S2**], we did not pursue this further: given the anatomical distributions of CO in lateral and ventral-only locations, it is hard to conceive a way in which they could detect dorsal-only contact over ventral-only [Hartenstein, 1988]; also, their known functions as proprioceptors [Caldwell et al., 2003] likely preclude a role in regional substrate detection.

In all cases, we observed little effect of temperature on larvae expressing *UAS-shi^[ts]^* in the absence of Gal4. Although there were some trends for shorter self-righting times at the restrictive temperature, this is in line with the known effects of temperature on other behaviours like crawling [Evans et al., 2023]. Overall, through conditional inhibition, these results demonstrate the importance of the proper function of multidendritic neurons for the triggering of an effective self-righting behaviour.

### Optogenetic inhibition of anterior but not posterior multidendritic neurons delays self-righting

Having observed that normal activity of multidendritic sensory neurons was important for self-righting, we wished to reconcile this finding with the previous observation of an anterior dominance in self-righting. Specifically, we aimed to determine if the anterior multidendritic neurons were responsible for registering the sensory inputs required to induce self-righting. To this end, we developed a novel approach to achieve spatially-restricted neuronal inhibition in a freely moving larva, a technique that we coined the “opto-axial method”. We designed a custom 3D-printed arena containing a small slit, which acts as a photomask for an LED positioned underneath [**Figure 4A**] This approach allowed us to target three segments on a third-instar larva, which we accurately positioned and recorded using the water-unlocking technique [**Figure 4B**]. To test the effect of neuronal location on self-righting, we placed each larva in five positions: four positions which targeted segments along the AP axis and one control position with no light [**Figure 4C**], following a randomised order. We elicited optogenetic inhibition of multidendritic neurons via expression of the photosensitive anion channel GtACR2 [Govnorova et al., 2014] in the 109(2)80-Gal4 domain.

**Figure 4:**
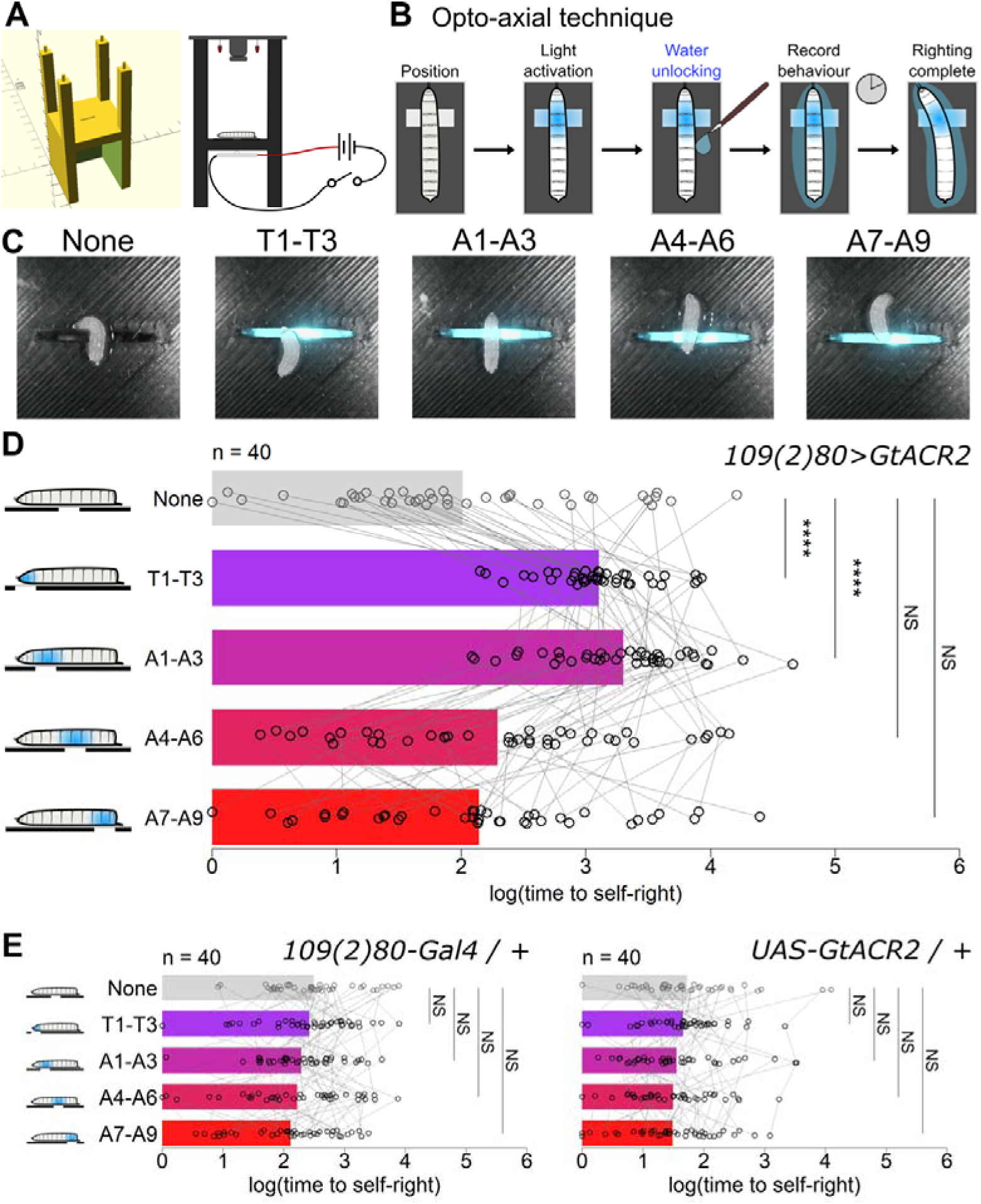
Localised optogenetic inhibition of multidendritic sensory neurons along the anterior-posterior axis. **A)** Experimental setup for localised optogenetic inhibition. Inhibitory light was spatially restricted by means of a small slit in the base of a 3D-printed arena (left). The LED was positioned under the slit and operated by a switch, while an infrared camera recorded from above (right). **B)** Experimental procedure for testing the effects of localised inhibition on self-righting. Larvae were positioned dorsal side down on a coverslip above the slit such that incoming light illuminated a group of three segments. After light activation, larvae were quickly unlocked through application of water and the time to complete self-righting was timed. **C)** Photographs of the five experimental conditions of localised illumination. In each condition, light targeted a group of three segments along the anterior-posterior axis, while no light was used as a control condition. **D)** Log-transformed time to self-right across the five illumination conditions for larvae expressing GtACR2 in the 109(2)80 domain. The bars show mean values, and points show individual measurements with measurements from the same larva being joined by grey lines. Analysis of deviance for a mixed model including control genotypes revealed a significant interaction of illumination condition and genotype (F_(8, 468)_ = 10.35, P < .001, n = 40). ‘****’ = P < .0001, NS = P > .05 for post-hoc one-sided comparisons between conditions of light illumination, following Dunnet’s approach with Sidak’s adjustment for multiple comparisons. **E)** Log-transformed time to self-right across the five illumination conditions for Gal4 (left) and UAS (right) genetic control lines. No statistical differences were observed between the control condition and the experimental illumination conditions for either control line.

Mirroring our results with substrate contact, we found that inhibition of multidendritic neurons in the anterior regions comprising segments T1-T3 and A1-A3 significantly prolonged self-righting compared to the no illumination condition. Conversely, inhibition in the more posterior regions comprising segments A4-A6 and A7-A9 had little effect on self-righting times [**Figure 4D**], suggesting that anterior multidendritic neurons are uniquely responsible for directing self-righting behaviour. Furthermore, the effect size of inhibition on self-righting time was similar for segments T1-T3 and A1-A3, indicating the existence of a relatively sharp transition in self-righting-relevant activity at around halfway along the AP axis, as opposed to a sensory gradient from posterior to anterior. We observed no differences in self-righting times between conditions of regional illumination in genetic control larvae that were subjected an identical experimental procedure **[Figure 4E]** showing this effect was indeed due to neuronal inhibition rather than light illumination alone. Visual inspection of recordings showed that while the no light and posterior conditions produced a relatively normal self-righting sequence in experimental larvae, the anterior inhibition appeared to lead to a more disorganised movement sequence with tight muscle contractions as has been reported previously for multidendritic inhibition [Hughes & Thomas, 2007]. In conclusion, these results demonstrate that specific activity of the anterior multidendritic neurons is key to effective and coordinated self-righting behaviour.

Given the current results of thermogenetic inhibition and those from prior studies showing a role of the daIV subpopulation in self-righting [Klann et al., 2021], we elected to repeat the opto-axial approach using the *ppk-Gal4* driver, which drives expression solely in daIV md neurons. We observed the same pattern as with the 109(2)80-Gal4 domain, with inhibition of daIVs in T1-T3 and A1-A3 regions leading to significantly longer self-righting times compared to no illumination, while inhibition in A4-A6 and A7-A9 had no effect. No changes in self-righting times according to illumination condition were observed in genetic control lines **[Figure S3]**. However, as with thermogenetic inhibition, the effect size of the self-righting delay was reduced compared to the 109(2)80-Gal4 domain. One possible explanation for this consistent difference in effect size is the relative expression strengths of the two Gal4 drivers. However, measurements of tdTomato fluorescence expressed in daIV neurons via the two drivers indicated no difference in expression level **[Figure S3C]**. This suggests that suggesting that md neurons other than daIVs play a role in facilitating the efficient completion of self-righting behaviour. Given that daIII neurons appeared not to contribute to normal self-righting, the daI or daII neurons could be involved, whose contribution may be more proprioceptive in nature.

### Inhibition of anterior multidendritic neurons is associated with increased head casting during self-righting

Although inhibition of anterior multidendritic neurons led to impaired self-righting behaviour, we were still unsure as to what role these localised neurons played in shaping the self-righting sequence. While visual examination of larval behaviour suggested that anterior sensory inhibition produced a more disorganised self-righting sequence, we questioned exactly how the self-righting sequence was changed. In other words, we wished to extract the precise features of movement that were altered under anterior inhibition using our framework for self-righting quantification first detailed in Figure 1. To obtain this quantification, we trained a separate network through DeepLabCut using recordings of larvae undergoing optogenetic inhibition that were trimmed to include only the behavioural sequence [**Figure 5A**]. As before, we achieved accurate tracking of four points along the AP axis that labelled the head, tail and two middle points.

**Figure 5:**
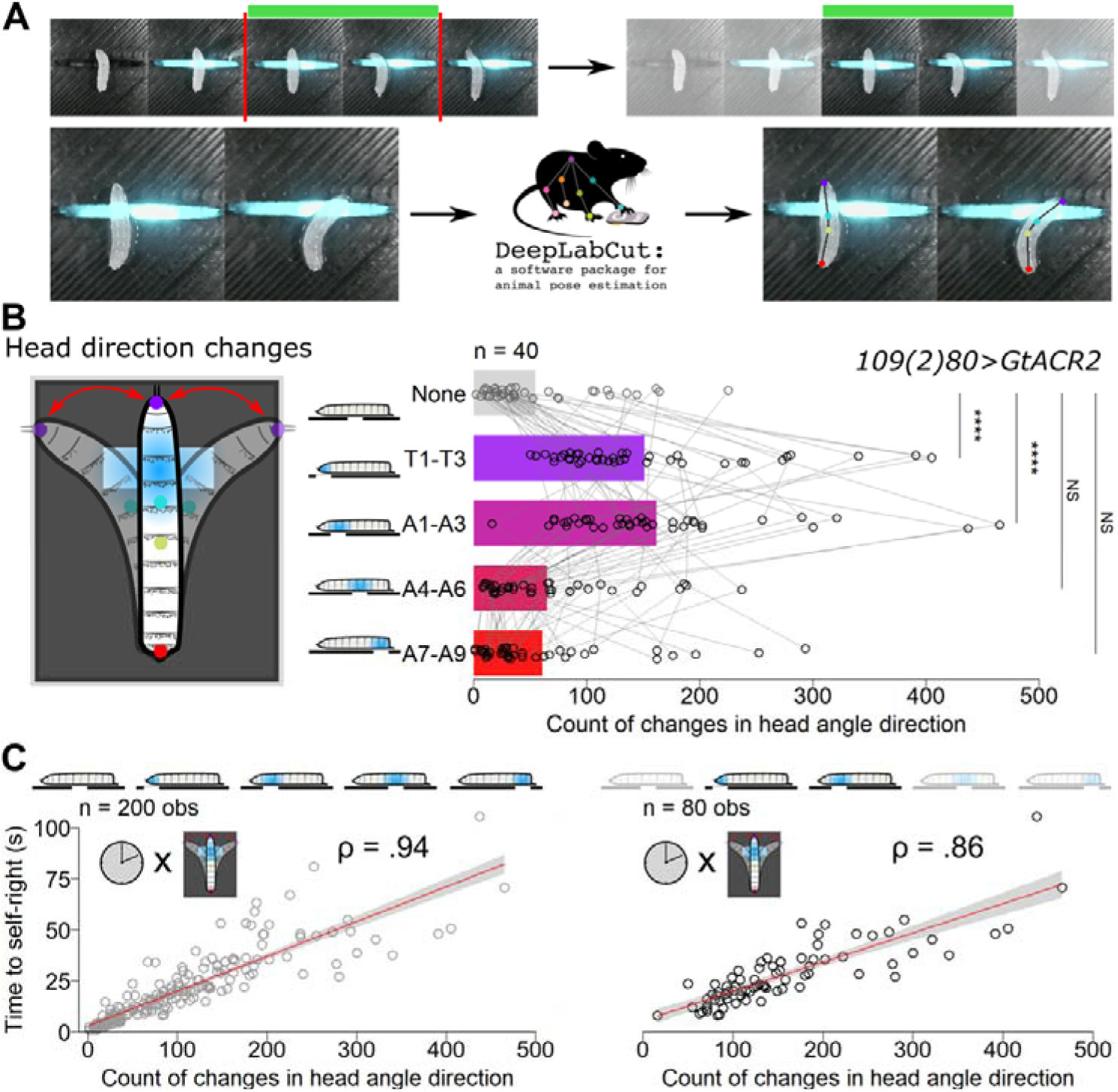
Behavioural changes occurring under localised optogenetic inhibition of sensory neurons. **A)** Labelling of recordings using DeepLabCut. Recordings of optogenetically-inhibited larvae were first manually trimmed so they contained only the time from movement unlocking to completion of self-righting (top). These videos were then analysed by DeepLabCut, which tracked four points along the anterior-posterior axis: head (purple), anterior middle (cyan), posterior middle (yellow), and tail (red). The coordinates of these tracked points were then used to calculate features of self-righting. **B)** Counts of the changes in head curvature direction under localised optogenetic inhibition of multidendritic neurons. The counts were calculated as the number of times a larva went from bending towards one direction with the head to bending in the other direction. Bars show the mean values for each condition, while points show counts for individual samples with measurements from the same larva being connected by lines. Analysis of deviance for a negative binomial model including control genotypes revealed a significant interaction of illumination condition and genotype (Χ^2^_(8)_ = 40.41, P < .001, n = 40). ‘****’ = P < .0001 for post-hoc one-sided comparisons between conditions of light illumination, following Dunnet’s approach with Sidak’s adjustment for multiple comparisons. **C)** Correlations between count of head direction changes and self-righting times, for all illumination conditions (left) and just the anterior illumination conditions (right) in 109(2)80>GtACR2 larvae. Points show individual observations, while the red line indicates a linear regression. P < .0001 for both Spearman correlations.

As our initial characterisation of self-righting behaviour revealed the importance differences between mean head and tail speeds, we again analysed this feature in larvae that underwent the regional optogenetic inhibition. For head speed, we observed a reduction in mean speed for experimental larvae, but only in the A1-A3 condition. Conversely, for tail speed, we observed reductions in both the T1-T3 and A1-A3 conditions [supplementary material **Figure S4**]. However, from this result alone it was still not entirely clear whether decreased speed was in fact related to increased self-righting times. To test the relationship between the two features, we calculated the spearman correlations between counts of mean head and tail speeds and self-righting times. For head speed, there was a moderate negative correlation with self-righting time for the 109(2)80>GtACR2 larvae across all illumination conditions (ρ = −.60, P < .001). Similarly for tail speed, there was a large negative correlation with self-righting time in the same conditions (ρ = −.73, P < .001). However, when we constrained the analysis to just the T1-T3 and A1-A3 conditions where self-righting time was experimentally delayed, the correlation magnitude dropped substantially for both head (ρ = −.16, P = .0159) and tail speeds (ρ = −.30, P = .007). This suggests that reductions in speed likely do not fully explain reductions in self-righting time caused by inhibition of anterior sensory neurons. Furthermore, we also observed an increase in head speed in the T1-T3 condition for UAS-GtARCR2 control larvae [supplementary material **Figure S4**]. These findings led us to infer that speed was not an ideal predictor for changes in self-righting time due to sensory manipulation and prompted us to search for other factors that were more strongly associated with self-righting delays. Given the importance of head and tail bending during self-righting behaviour, we reasoned that the degree of curvature could be altered across conditions of regional illumination. However, we found no differences in mean absolute curvature across any illumination conditions for any genotypes [supplementary material **Figure S5**], suggesting that the ability of larvae to form bends appropriate to self-righting behaviour was not impacted by our experimental manipulation. Therefore, it appeared self-righting delays under anterior sensory inhibition were not due to larvae simply moving more slowly than normal or not achieving the optimal degree of curvature required for self-righting.

We reasoned that instead of simple changes to high-level features of larval movement, anterior sensory inhibition may be impacting the precise pattern of movements prior to self-righting completion. Another visual inspection of recordings showed that while head speed and curvature was indeed comparable to the normal sequence, the head appeared to make many more back-and-forth movements under anterior sensory inhibition, reminiscent of head casting behaviour [Ohyama et al., 2013]. We sought to quantify this observation by counting the number of times that larvae changed the head angle direction; that is, going from bending towards the right to bending towards the left or vice versa. In comparing the count of head direction changes across the regions optogenetic inhibition, we observed a pattern that strikingly matched that of the self-righting times, with significantly increased counts in the anterior T1-T3 and A1-A3 conditions but not the posterior A4-A6 and A7-A9 conditions [**Figure 5B**]. We also observed no increases in counts of head curvature changes in genetic control larvae [supplementary material **Figure S6**]. This result confirms our observation that inhibition of anterior multidendritic neurons causes a shift to a more disorganised self-righting sequence that is characterised by a high preponderance of head casting behaviour. This finding could reflect the possibility that under anterior sensory inhibition larvae are unable to properly detect the underlying surface, and so employ a repeated head sweeping behaviour in order to find a suitable substrate for attachment.

As we did previously for movement speeds, we wanted to know if this increased head casting behaviour was relevant to increases in self-righting times. We first calculated spearman correlation coefficients for 109(2)80>GtACR2 larvae across all illumination conditions, to see if number of head casts was generally related to differences in self-righting times. Here, we found an extremely strong correlation (ρ = .94), showing that the natural variability in the timing of the self-righting sequence can be largely accounted for by the number of head casts. However, we also wished to see if this relationship held in the experimental conditions of anterior sensory inhibition where self-righting time was prolonged. When we restricted data to the T1-T3 and A1-A3 conditions, we observed a slightly reduced albeit very strong correlation (ρ = .86) between head direction changes and self-righting times [**Figure 5C**]. This supports the notion that this natural relationship still holds for experimental conditions of anterior sensory perturbation, where larvae performed much more head casting overall. In conclusion, our quantification of the features of self-righting shows that anterior sensory inhibition does not alter simple features of speed or curvature but rather causes a behavioural switch to increased head casting behaviour that well explains the delays in self-righting completion.

Since inhibition of the anterior daIVs specifically also resulted in increases in self-righting times, we questioned if an increase in head casting behaviour also occurred with this sensory subpopulation. Indeed, we observed significantly increased number of head casts in the T1-T3 and A1-A3 conditions, but not in the A4-A6 or A7-A9 conditions. As with all md neurons, this the number of head direction changes was significantly correlated with self-righting times, across both all conditions and the anterior illumination conditions only **[Figure S7]**. This suggests that the switch to head casting behaviour over self-righting under anterior sensory inhibition is unlikely to be driven primarily by impaired proprioceptive input, but rather by an inability to sense the underlying substrate. We speculate that increased head casting under anterior sensory inhibition likely reflects a search strategy for a suitable substrate, as the larvae are unable to detect the substrate underneath them on which they would typically self-right.

### Normal expression of the Drosophila *Hox* genes *Antennapedia* and *Abdominal-B* in multidendritic neurons is required for self-righting behaviour

Our results thus far demonstrated a clear anterior precedence of self-righting behaviour, which involves not only anterior-dominated movements but also a key role of the anterior sensory system in detecting the substrate contact that elicits this postural behaviour. We thus wondered what genetic mechanisms may underlie the establishment of this pattern. Given the clear discrepancies of self-righting behaviour aligning with the AP axis, we questioned if the Hox genes, which play a key role in the developmental allocation of cellular identity along to this very axis, might play a role.

To first address this possibility, we first sought to determine if the sensory neurons involved in self-righting did indeed express any Hox genes at the larval stage. For this we used fluorescence-activated cell sorting (FACS) to collect multidendritic neurons from first instar larvae expressing GFP in the 109(2)80-Gal4 domain, extracting RNA from collected cells to carry out RT-PCR to amplify products derived from the four Hox genes: *Antennapedia* (*Antp*), *Ultrabithorax* (*Ubx*), *abdominal-A* (*abd-A*) and *Abdominal-B* (*Abd-B*). We selected these Hox genes based on their axial expression domains across other, relevant tissues (epidermis, central nervous system) of the embryonic and larval *Drosophila* [Mallo & Alonso, 2013]. Analysis of the resulting amplicons confirmed expression of all *Hox* genes tested; however, bands were particularly bright and sharp for *Antp* and *Abd-b* [**Figure 6A**], indicative of strong expression of *Antp* and *Abd-b* RNA in multidendritic neurons.

**Figure 6:**
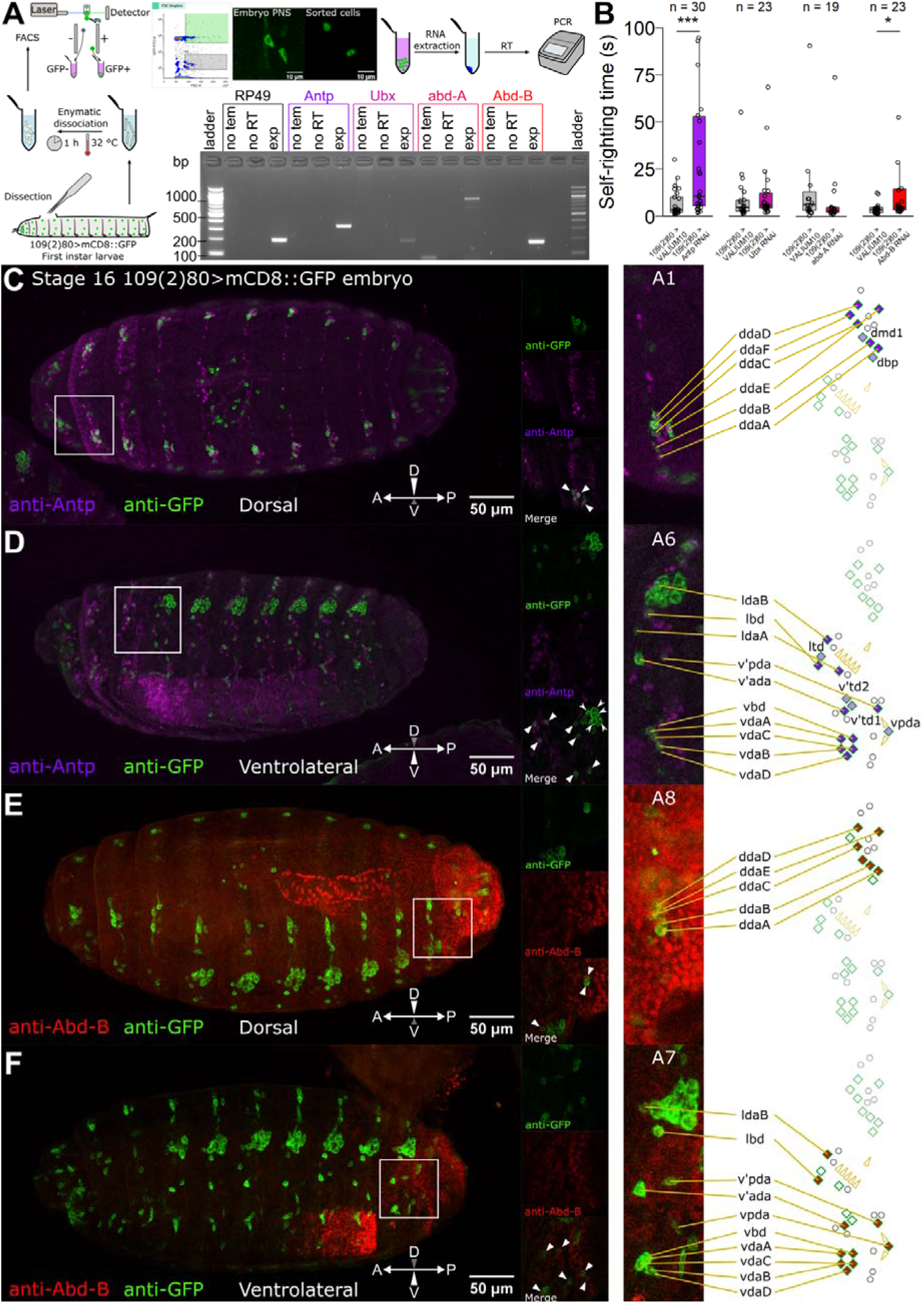
Hox expression in the sensory system and its influence on self-righting behaviour. **A)** Quantification of Hox RNA expression in larval sensory neurons. The flow diagram (beginning bottom left) shows how sensory neurons were collected from dissected first instar larvae using fluorescence-activated cell sorting (FACS). RNA was extracted from the sorted cells and reverse transcribed to DNA for amplification of Hox gene products via PCR. The photograph (bottom right) shows an agarose gel electrophoresis following RT-PCR of Hox genes *Antp*, *Ubx*, *abd-A* and *Abd-B*. For each gene, ‘no tem’ indicates a no template cDNA control, ‘no RT’ indicates a no reverse transcription control, and ‘exp’ indicates the experimental lane. **B)** Self-righting times of first instar larvae expressing a Hox gene RNAi construct in the 109(2)80 domain. ‘***’ = P < .001, ‘*’ = P < .05 for Wilcoxon rank sum tests, n = 19-30. **C-F)** Confocal images of immunolabelled stage 16 109(2)80>mCD8::GFP embryos. In each case, the large left panel is a maximum intensity z-projection. The white square indicates the region that is zoomed in the smaller panels to the right. These smaller panels show an individual z slice in the two separate channels and the channel overlay. Triangles indicate cells showing clear signal for Hox protein, and arrowheads indicate cells lacking signal for Hox protein. The strips on the right are zoomed sections of the z-projection, showing a range of cells in one embryonic hemisegment. The putative identities of these cells are indicated by red lines in accordance with the canonical hemisegmental diagram on the far right. **C)** Embryo immunolabelled for Antp and GFP, dorsal view. **D)** Embryo immunolabelled for Antp and GFP, ventrolateral view. **E)** Embryo immunolabelled for Abd-B and GFP, dorsal view. **F)** Embryo labelled for Abd-B and GFP, ventrolateral view.

Having observed Hox expression in larval multidendritic neurons, we next questioned if a specific level of Hox expression was important for sensory contributions to self-righting behaviour. We hypothesised that expression of the more anterior Hox genes – *Antp* and *Ubx* – would influence self-righting, given the importance of anterior sensory neurons in self-righting. To test this hypothesis, we used RNAi to knockdown expression of the above noted four genes within the *109(2)80-Gal4* domain of first instar larvae, and tested self-righting times against an empty RNAi control. Intriguingly, while RNAi against *Antp* induced a substantial delay of self-righting times, no effect was observed for *Ubx*; instead, we found RNAi against *Abd-b* also produced a statistically significant, yet modest, increase in self-righting times [**Figure 6B**]. This latter finding may be related to the known role of posterior segments in guiding motor sequences, as has been recently observed for locomotion [Jonaitis et al., 2024]. Conversely, the former finding could be explained by Antp expression within anterior sensory neurons being essential for normal activity levels, as has also been demonstrated for Hox genes in other cellular populations [Picao-Osorio et al., 2015; Issa et al., 2019]. Given our previous results regarding daIV neurons, we also questioned if knockdown of Hox genes within this population also resulted in self-righting delays. Interestingly, we observed no effect of Hox gene knockdown within the ppk-Gal4 domain [**Figure S8**], suggesting that multidendritic neurons other than daIVs are subject to Hox effects that ultimately impact self-righting.

However, the observation of Hox RNA expression in the multidendritic population does not necessarily imply Hox protein translation. Furthermore, we lacked information about the exact neurons in which *Antp* and *Abd-b* could be exerting their effects on self-righting, information that should aid our understanding of the sensory processing underlying self-righting. To address these open aspects, we carried out a series of fluorescence immunolabelling experiments to map the expression of Antp and Abd-B proteins in late-stage *109(2)80>mCD8::GFP* embryos to determine whether these proteins were expressed in the multidendritic population. As we had discovered that self-righting behaviour relies on precise mechanical stimulation at different regions of the body, we endeavoured to capture expression in the different clusters of the developing PNS. For both Hox genes, we observed the expected pattern of expression within the developing CNS, with Antp showing high expression in the anterior of the VNC [**Figure 6D**] and Abd-b displaying strong signal at the very posterior of the VNC [**Figure 6F**], providing a natural internal control for our experiments.

Analysis of the PNS pattern for Antp showed, unexpectedly, signal throughout the entire PNS, in the dorsal [**Figure 6C**] as well as the ventral and lateral clusters of multidendritic neurons [**Figure 6D**]. However, the intensity of the observed signal was suggestive of a trend with higher values in the anterior. With regard to specific neuronal classes, examination across z slices revealed that Antp was expressed in the majority of neurons labelled by 109(2)80-Gal4, save for the lateral cluster of oenocytes that also fall into this domain [Grueber et al., 2002; Klann et al., 2021]. Additionally, we were unable to precisely confirm Antp expression within the tracheal branching and bipolar multidendritic neurons, which could be due to the reduced expression strength of these classes in the 109(2)80-Gal4 domain compared to the dendritic arborisation neurons [Hughes & Thomas, 2007]. These observations lead us to conclude that Antp is likely expressed within all dendritic arborisation neurons of the PNS with a potentially modestly higher level in anterior regions.

Conversely, the expression of Abd-b within the PNS was more limited. While we observed expression in the dorsal [**Figure 6E**], ventral and lateral clusters [**Figure 6F**], strong signal was restricted to segments A7 and A8, mirroring the pattern of the CNS. As with Antp, Abd-B signal was observed in most neurons of those segments, suggesting a role for Abd-B in the posterior-most dendritic arborisation neurons. However, due to the restricted expression domain, as well as the uncharacterised nature of the multidendritic neurons in A8, we cannot assert that Abd-B is expressed in all the dendritic arborisation neurons of these two segments. Regardless, given our findings that self-righting relies on sensory information from the anterior, we suggest that the role of Abd-B in this behaviour is likely to be attributed to the regulation of motor patterns through proprioceptive feedback. Taken together, these results show that at least two Hox genes – *Antp and Abd-b* – are expressed within sensory neurons of the late embryo and early larva, and that this expression is key for optimal sensory function in the context of self-righting behaviour. That these two Hox genes, but not *Ubx* or *abd-A*, affect self-righting via the sensory system might suggest a role of the two termini of the animal in interpreting sensory information for complex behaviours like self-righting.

## DISCUSSION

### Mapping the behavioural effects of regional stimulation in Drosophila larvae

The interplay between mechanical stimuli, sensory neuron activity and specific behaviours has been well-studied, with Drosophila research leading progress in this field [Hughes & Thomas, 2007; Ohyama et al., 2013; Titlow et al., 2014; Ohyama et al., 2015; Jovanic et al., 2016; Burgos et al., 2018; Liu et al., 2022]. Here we extend this work by examining the influences of body plan, morphology, and axial patterning on the relationship between sensory function and behaviour. Through examination of postural control, as instantiated in the highly conserved self-righting behaviour, we sought to address how localised sensory function relates to the coordinated movement of specific body parts. Using a novel manipulation technique that we coin ‘water-unlocking’, we first characterised the self-righting sequence as involving a strong movement and curvature of the anterior, which increases in magnitude over the course of the behaviour. To test how this behaviour is related to sensory contexts, we restricted substrate contact to different regions of the body, finding that anterior substrate contact on the dorsal side is required for self-righting while anterior contact at the ventral side prevents its occurrence.

Wishing to know which sensory neurons underlie this anterior precedence, we assayed self-righting under conditional inhibition of subpopulations of sensory neurons. We observed that inhibition of multidendritic neurons, in particular that of daIV neurons, led to deficits in self-righting behaviour, in agreement with previous results employing constitutive inhibition [Klann et al., 2021]. However, it remained unclear how the localised activity of these neurons might shape behavioural responses. To address this issue, we developed a new technique that we term the ‘opto-axial approach’, which involved optogenetic inhibition of groups of sensory neurons along the AP axis [Video S1]. In alignment with our previous results, we found that inhibition of multidendritic neurons in the thoracic and anterior abdominal segments led to strong deficits in self-righting behaviour, while inhibition at posterior segments had no discernible effect. These results confirm a role of anterior multidendritic activity in driving the head-dominated self-righting behaviour, and agree with prior studies observing a particular sensory sensitivity of the anterior body [Murawski et al., 2020]. However, although anterior sensory function appeared to be important for self-righting, it was not clear how disrupting this activity was affecting the behavioural sequence. Using automated tracking [Video S2] of body parts along the AP axis, we found that inhibition of anterior sensory neurons produced a unique over-representation of head casting behaviour, which was strongly correlated with delays in time taken to complete self-righting. This suggests that despite the important role of head movement in self-righting, a lack of normal sensory activity at the anterior causes a switch to different behavioural sequence characterised by excessive head movement that fails to convert to the rest of the behavioural sequence. The same patterns of self-righting delays and behavioural changes were also observed for axial inhibition of daIV neurons specifically, suggesting this subpopulation of md neurons are important for inferring postural state. However, stronger effect sizes for inhibition of the whole md population also indicates that other subpopulations of the md neurons, such as the daIs or daIIs, contribute to normal self-righting completion. We speculate that given their wide dendritic arbors and ability to detect externally applied pressure [Liu et al., 2022], the daIV neurons most likely detect substrate contact and allow the inference of position with respect to external surfaces. In contrast, the daI neurons, which have a known role in proprioception [Vaadia et al., 2019], likely detect the body configuration such as the magnitude of bends and torsion. The combination of these different somatosensory elements is likely necessary for a typical and efficient self-righting sequence.

Finally, we questioned which genetic mechanisms might contribute to the observed patterns of anterior sensory dominance. Given the discrepancies of activity according to the AP axis, we hypothesised that the Hox genes may play a role. Through a combination of cell sorting, PCR, immunolabelling and RNAi experiments, we found that the Hox genes *Antp* and *Abd-b* are expressed in different subsets of the multidendritic sensory neurons and that typical expression levels are required for normal self-righting behaviour. Altogether, we show that the correspondence of mechanosensory stimuli and postural behaviour occurs via region-specific mechanisms that are tightly linked to body plan and its underlying genetic specification.

### Exploring the relationship between body form and function via sensory biology

Our finding that anterior sensory activity is required for postural control behaviour highlights the intimate relationship between body form and function. Previous research in Drosophila has primarily focused on how behaviours relate to activity in different sensory cell types [Hughes & Thomas, 2007; Burgos et al., 2018; Vaadia et al., 2019]. These cell types arise from differing developmental lineages and possess varying morphologies [Grueber et al., 2002; Singhania & Gruber, 2014], making them ideal candidates for detecting stimuli of differing modality and intensity. However, this line of investigation, though insightful, does not fully integrate the notion that behaviour is enacted through the spatial form of the body and thus also depends on the spatial configuration of the environmental stimuli that induce it.

Previous research indicated that larval responses to the same mechanosensory stimulus do depend on its location of delivery to the body [Titlow et al., 2014; Murawski et al., 2020], as well as other contextual factors such as co-occurring stimuli [Ohyama et al., 2015; Jovanic et al., 2016]. Neuronal manipulations have further shown that these differing responses, such as the directionality of crawling in response to a pin prick at the head or tail, depend on the localised activity of segmentally repeated interneurons that preferentially feed activity into neighbouring motor pattern circuits [Takagi et al., 2017]. Our work extends this line of investigation by characterising the role of regional sensory components of the same cell type, enabling us to tease apart cell type specification roles from regional specific roles. In our understanding, we show for the first time that a complex behavioural sequence varies greatly according to the precise location of sensory activity. Specifically, we demonstrate that activity of anterior sensory neurons is key for the Drosophila larva to correct its posture with regard to the underlying substrate. That this self-righting behaviour also involves predominant movements of the head may highlight a general principle for the coordination between sensory activity and related muscle activations, indicating a possible regional, modular specialisation of neuronal circuits to reduce wiring cost and maximise efficient information flow [Rubinov, 2016; Goulas et al., 2019].

Furthermore, our finding that inhibition of anterior sensory activity does not eliminate behaviour, but instead causes a switch to increased head casting, also reflects the intimate relationship between body plan and behaviour. Despite presumably normal posterior sensory activity – and capable musculature – larvae still preferentially move with the head, a pattern we also observed during posterior-only substrate contact. We suggest this is due to the anterior dominance also observed during crawling behaviour, which is initiated with a forward grabbing with the mouth hooks [Liu et al., 2023], and which we also observed to occur under anterior ventral contact. Following from these observations, our results reflect the general principle that the sensory capacity of animals is reciprocally constrained by their behavioural repertoire, which is, in turn, further constrained by their spatial form [Salinas, 2006; Lungarella & Sporns, 2006; Molano-Mazon et al., 2023]. This goes against the notion of a one-to-one correspondence between sensory stimuli and behaviours, and, instead, points to a model in which movements are dynamically shaped by a multitude of contextual factors with spatial location being a key driver of behavioural diversity. These findings bear implications for future studies of sensory function and behaviour, since properties of a specific cell type, such as neuronal activity and synaptic connectivity, may well differ according to body localisation. Indeed, it remains an open question as to how mechanical stimuli at different locations can produce entirely different behavioural sequences. The recent ultrastructural mapping of sensory sensilla in the Drosophila larva [Richter et al. 2025] should pave the way to further investigations on how local stimulations trigger the complex repertoire of larval behaviours.

### A novel role of the Hox genes in Drosophila SNs

Our finding that Hox gene expression within sensory neurons is required for normal self-righting behaviour presents a genetic mechanism by which the properties of the PNS are shaped according to position along the AP axis. In the larval CNS, expression of individual Hox proteins is constrained to localised domains along the AP axis, where they act to specify neuroblast lineages that eventually communicate with corresponding body parts in the head, thorax and abdomen [Prokop et al., 1998; Technau et al., 2006; Becker et al., 2016]. Hox expression has also been shown to be important for the specification of components of the PNS, including sensory organ precursor (SOP) formation [Gutzwiller et al., 2010] and PNS structure [Mann & Hogness, 1990; Heuer & Kaufman, 1992]. However, our work shows – in our understanding – for the first time that Hox expression is important for the normal function of the multidendritic sensory neurons. Accumulating evidence from studies in the CNS suggests that in addition to their roles in developmental specification, Hox genes also regulate post-developmental neuronal functions [Picao-Osorio 2015, Issa et al 2019, Issa et al. 2022]. For example, recent research has demonstrated that self-righting behaviour in the adult fly depends on ongoing expression of *Ubx* in NB2-3/lin15 motor neurons, which regulates plastic connectivity between these neurons and their target muscles [Issa et al., 2019]. At the larval stage, post-developmental upregulation of *Ubx* in a subset of lateral transverse (LT) motor neurons is sufficient to perturb self-righting behaviour [Picao-Osorio et al., 2015]. Based on this work, it is possible to conceive that the effects of Hox disruption we observe in the PNS could be due to a developmental effect on multidendritic neurons, an effect on neuronal physiology, or a combination of both. Indeed, previous work has shown that perturbations to Hox expression, either directly or through manipulation of Polycomb-group proteins, has a substantial impact on the dendritic structure of the da neurons. Specifically, overexpression of several Hox genes including Ubx and Abd-A leads to under-developed dendritic arborisations, while Hox loss of function mutations promote lability of terminal dendritic branches; notably, this regulation occurs post-mitotically, suggesting an ongoing role of dendrite maintenance rather than a developmental specification [Parrish et al., 2007]. More recent work also shows that the dendritic pruning of da neurons which occurs during metamorphosis is regulated by Hox genes, with overexpression of several Hox genes including Ubx, Abd-B and Scr leading to reduced pruning [Bu et al., 2023]. Therefore, Hox expression within the da neurons is likely to regulate dendrite morphology. Interestingly, despite some of these results being specific to daIV neurons, we found no difference in self-righting times when Hox knockdown was restricted to the daIV population. This indicates that md neurons other than daIVs are subject to Hox effects that are important for self-righting. Following the results from prior work, Hox knockdown within md neurons could increase dendritic branching and thus also the sensitivity of md neurons, possibly leading to improper action selection in response to dorsal substrate contact. Visual examination of recordings suggests that in contrast to the opto-axial experiments where larvae displayed head casting, the larvae expressing Hox RNAi in md neurons showed no apparent increase in head casting but instead performed continuous forwards and backwards peristaltic waves. Speculatively, given the lack of effect in daIV neurons, this could arise from overactive daI, daII or daIII populations, leading to the selection of crawling behaviour rather than self-righting.

Despite observing expression of several Hox genes (*Antp, Ubx, abd-A and Abd-B*) in multidendritic neurons, behavioural phenotypes were only observed for *Antp* and *Abd-B* knockdowns. This is especially interesting given the prior findings of the role of Ubx in dendritic arborisation of da neurons [Parrish et al., 2007], and could suggest differential importance of Hox genes for various behaviours according to their spatial domains and the neurons involved. Interestingly, while we hypothesised that *Antp* would be restricted to thoracic multidendritic neurons, as is observed in the CNS [Rosales-Vega et al., 2023], we observed *Antp* expression in multidendritic neurons along the entire AP axis. While the regulatory sequences controlling *Antp* expression in the PNS have been previously characterised [Boulet & Scott, 1988], we show conclusively that *Antp* expression can extend beyond its traditionally ascribed transcriptional domain well into the entire embryonic PNS. The strong effects of *Antp* disruption on self-righting behaviour could therefore reflect an important regulatory role for *Antp* in general sensory function, which we are currently investigating. In contrast, the expression of *Abd-B* followed the expected pattern, with expression observed only in the posterior-most multidendritic neurons. Given the anterior dominance of self-righting, it is somewhat unexpected that expression changes in posterior sensory neurons could perturb the behaviour. One possibility to explain this effect is that *Abd-B* knockdown causes a misinterpretation of mechanosensory signals in posterior sensory neurons, causing a switch to a different behaviour triggered by posterior stimulation. However, given the relatively small effect of *Abd-B* knockdown, we instead suggest that the posterior sensory neurons play a mostly proprioceptive role during self-righting behaviour, contributing to the stability of posterior segments during the head-dominated self-righting sequence. It remains an open question as to exactly how Hox expression impacts self-righting behaviour via regional sensory regulation. Aside from the known effects on da neuron dendritic morphology [Parrish et al., 2007], mechanisms could include the local specification of cell identity, synapse formation, and neuronal sensitivity via the regulation of channel proteins. For example, there is evidence for a role for the anterior *Hoxd1* expression in species-specific development of nociceptive connectivity in mouse and chick [Guo et al., 2010], posterior Hox gene expression in the identity of touch-responsive neurons in *C. elegans* [Zheng et al., 2015], and *Hoxa2* expression in the somatotopic organisation of the mouse brainstem [Bechara et al., 2015]. Therefore, it is possible that proper Hox expression in *Drosophila* multidendritic neurons affects their connectivity to downstream interneurons that direct activity towards specific motorneuron circuits. Ongoing work in our laboratory is seeking to address if Hox expression in the Drosophila PNS affects neuronal development, regional morphology, or physiology, as well as how this expression is important for the normal execution of the extensive repertoire of behaviours displayed by the fly larva and adult.

### Self-righting as an urbilaterian behaviour?

The present work has demonstrated that regional coupling between sensory function, body plan and movement is vital for an animal to properly orient itself within its environment. That righting and analogous behaviours are observed across bilaterian organisms, including invertebrates [Faisal & Matheson, 2001; Bilbao et al., 2018], fish [Bagnall & Schoppik, 2018], birds [Seal et al., 2019] and mammals [Jusufi et al., 2011; Feather-Schussler & Ferguson, 2016] raises the question of whether self-righting is an ancient behaviour. Indeed, the very nature of bilaterian organisms is such that the body is asymmetrically organised along the DV axis to facilitate specialised interactions with the environment. In this context, the ancestral bilaterian, the urbilaterian, has been suggested to have had a complex benthic adult form with AP and DV patterning, and that the key regulators of this patterning have been conserved throughout evolution [Robertis & Tejeda-Muñoz, 2022]. Given the requirement for a benthic adult to be anchored via its ventral side to the seabed, we speculate that self-righting behaviour may have been present in this ancient ancestor. In this context, self-righting could represent a fundamental movement that shaped body plans throughout bilaterian evolution, whereby the ability to actively invert the DV axis promoted further specialisation of the two sides of the body. While it is well accepted that the urbilaterian possessed an anterior primitive eye of some sort [Nilsson & Arendt, 2008], it is unclear to what degree other sensory *apparati* may have been specialised according to the AP axis. Future studies examining the relationship between patterning genes and PNS structure will be required to narrow down the postulated regional mechanosensory and neural specialisation of the urbilaterian.

## ACKNOWLEDGEMENTS

We wish to thank all members of the Alonso Lab for helpful discussions and feedback on this work. We would like to acknowledge our Sussex colleagues: Andre Chagas Maia for his advice and assistance with 3D printing of components, Yan Gu for expert support to our microscopy methods and Jonathan Wing for his help with FACS. We also wish to thank the Bloomington Drosophila Stock Center and the Vienna Drosophila Research Center for supplying many of the fly stocks used in this study. This research was funded by Leverhulme Trust Doctoral Scholarship given to WR, and a UK Medical Research Council Project Grant (MR/S011609/1) and a UK Biotechnology and Biological Sciences Research Council Project Grant (BB/Y006860/1) both awarded to CRA.

## Notes

### Competing Interest Statement

The authors have declared no competing interest.

### Summary of Updates

This revised version includes several new supplementary figures as well as it incorporates many changes in the text that improve the quality of the manuscript

