## Supplementary Materials for "Region-specific mechanosensation modulates *Drosophila* postural control behaviour"

*Department of Neuroscience,*

*Sussex Neuroscience,*

*University of Sussex,*

*Brighton BN1 9QG,*

*UK*

**This file includes:**

Figures S1 to S5

Legends for Movies 1 and 2

**Other supporting materials for this manuscript include the following:**

Movies 1 and 2

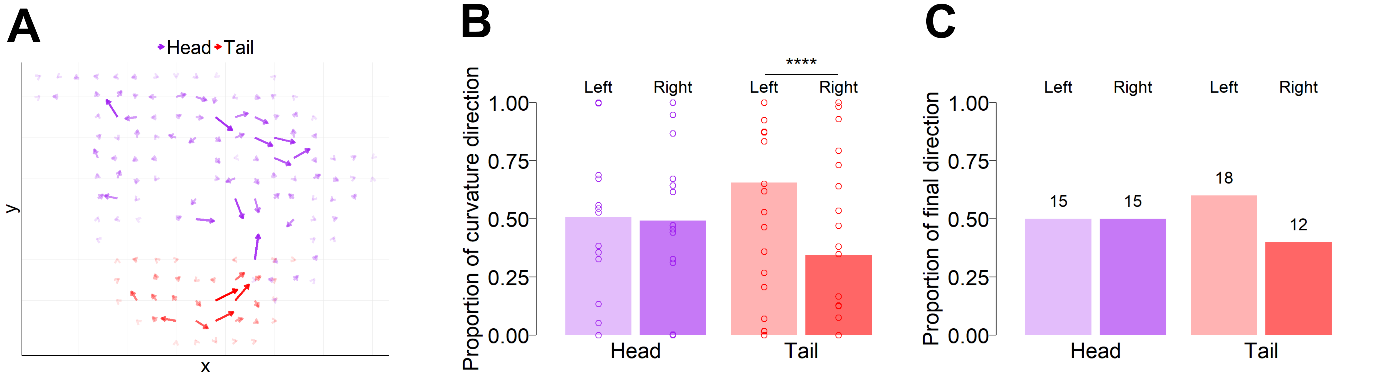

**Figure S1: Handedness of self-righting behaviour in 30 wild-type first instar larvae. A)** Vector field plot of head and tail movements during SR behaviour. Vectors were generated by rounding coordinate data to the nearest 10 pixels, then summing coordinate changes over the time course of the behaviour. The length and opacity of the arrows represents the magnitude of the vector sums, reflecting the tendency for larvae to move the head or tail in a given direction within the x-y plane. **B)** The proportion of time spent in left-handed or right-handed bends of the head or tail during SR behaviour. Bars indicate the mean proportion and points indicate proportions for individual larvae. ‘****’ = P < .0001 for a two-sided exact binomial test with a null probability of 0.5, n = 30. **C)** The proportion of larvae that completed the SR sequence with a left or right bend of the head and tail. Bars indicate the proportion, while the values above the bars indicate the number of larvae that displayed that direction of bend.

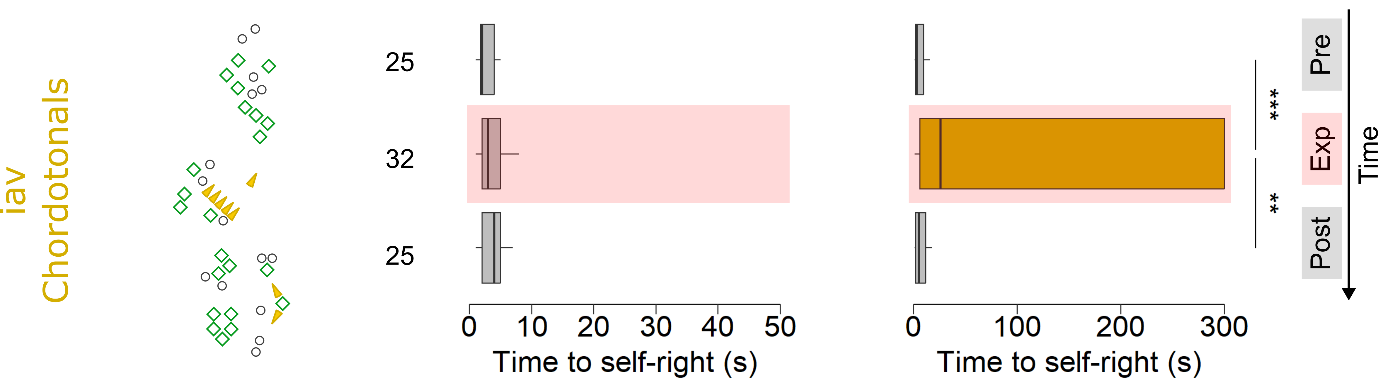

**Figure S2. Self-righting times for first-instar larvae expressing *shibire* in the chordotonal organs.** The location of the chordotonal organs within the hemisegmental sensory arrangement is shown (left). Box and whisker plots indicate self-righting times in each temperature condition, for *UAS-shi^[ts]^* controls (left) and age-matched experimental genotypes (right). The temporal order of the temperature conditions follows from top to bottom. ‘**’ = P < .01, ‘***’ = P < .001 for pairwise Wilcoxon signed-rank tests.

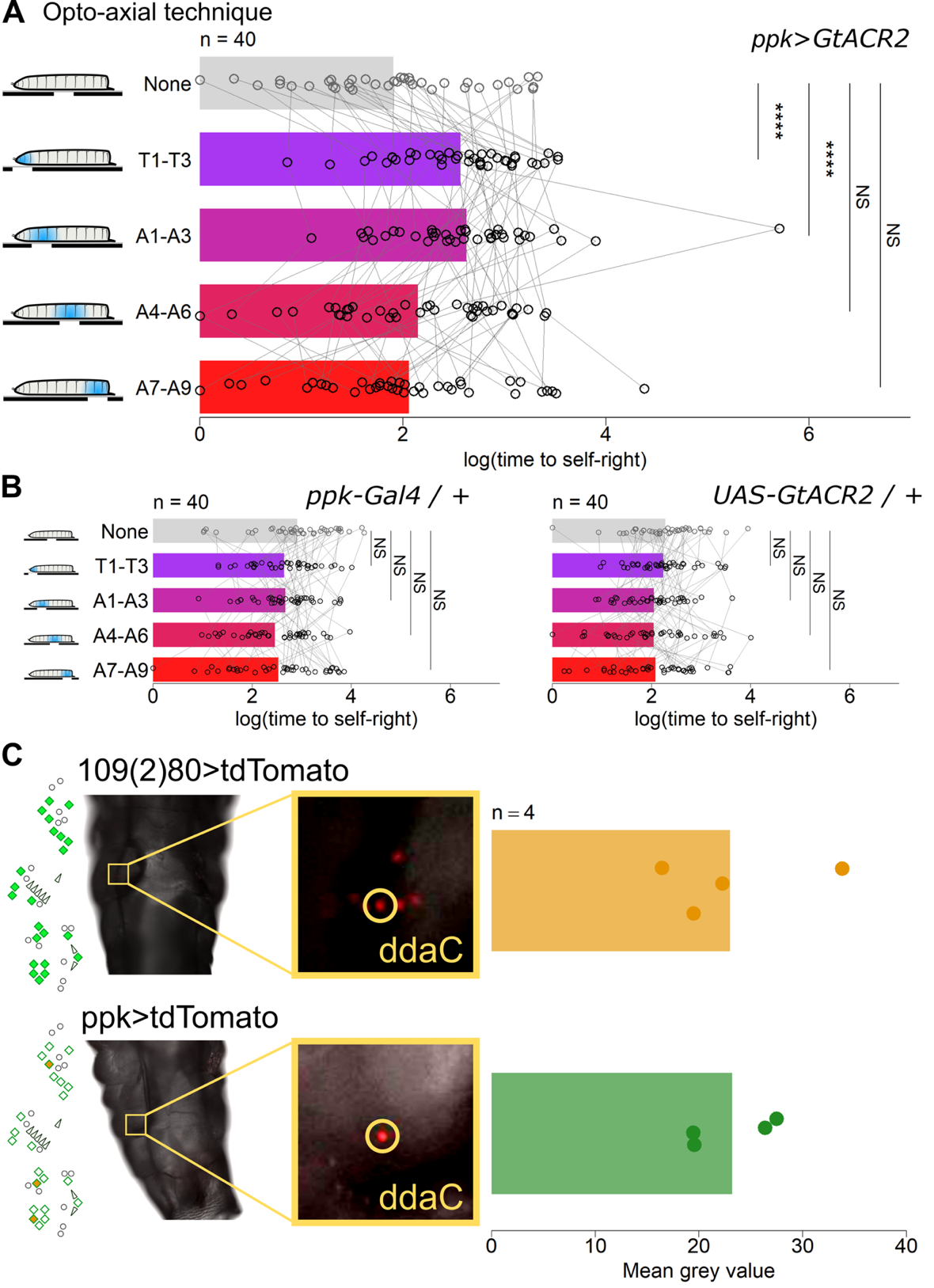

**Figure S3**. **Self-righting effects of localised optogenetic inhibition of daIV sensory neurons along the anterior-posterior axis. A)** Log-transformed time to self-right across the five illumination conditions for larvae expressing GtACR2 in the ppk domain. The bars show mean values, and points show individual measurements with measurements from the same larva being joined by grey lines. Analysis of deviance for a mixed model including control genotypes revealed a significant interaction of illumination condition and genotype (F_(8, 468)_ = 5.94, P < .001, n = 40). ‘****’ = P < .0001, NS = P > .05 for post-hoc one-sided comparisons between conditions of light illumination, following Dunnet’s approach with Sidak’s adjustment for multiple comparisons. Note that one single observation is not shown for visualisation purposes (value = 300 s). **B)** Log-transformed time to self-right across the five illumination conditions for Gal4 (left) and UAS (right) genetic control lines. No statistical differences were observed between the control condition and the experimental illumination conditions for either control line. **C)** tdTomato fluorescence intensity in ddaC expressed under 109(2)80 and ppk Gal4 drivers. The left panel shows a brightfield image of a third instar larva merged with the red fluorescence channel. The box shows a zoomed image of the region containing ddaC, which is circled. The right panel shows the mean grey value across four larvae expressing tdTomato.

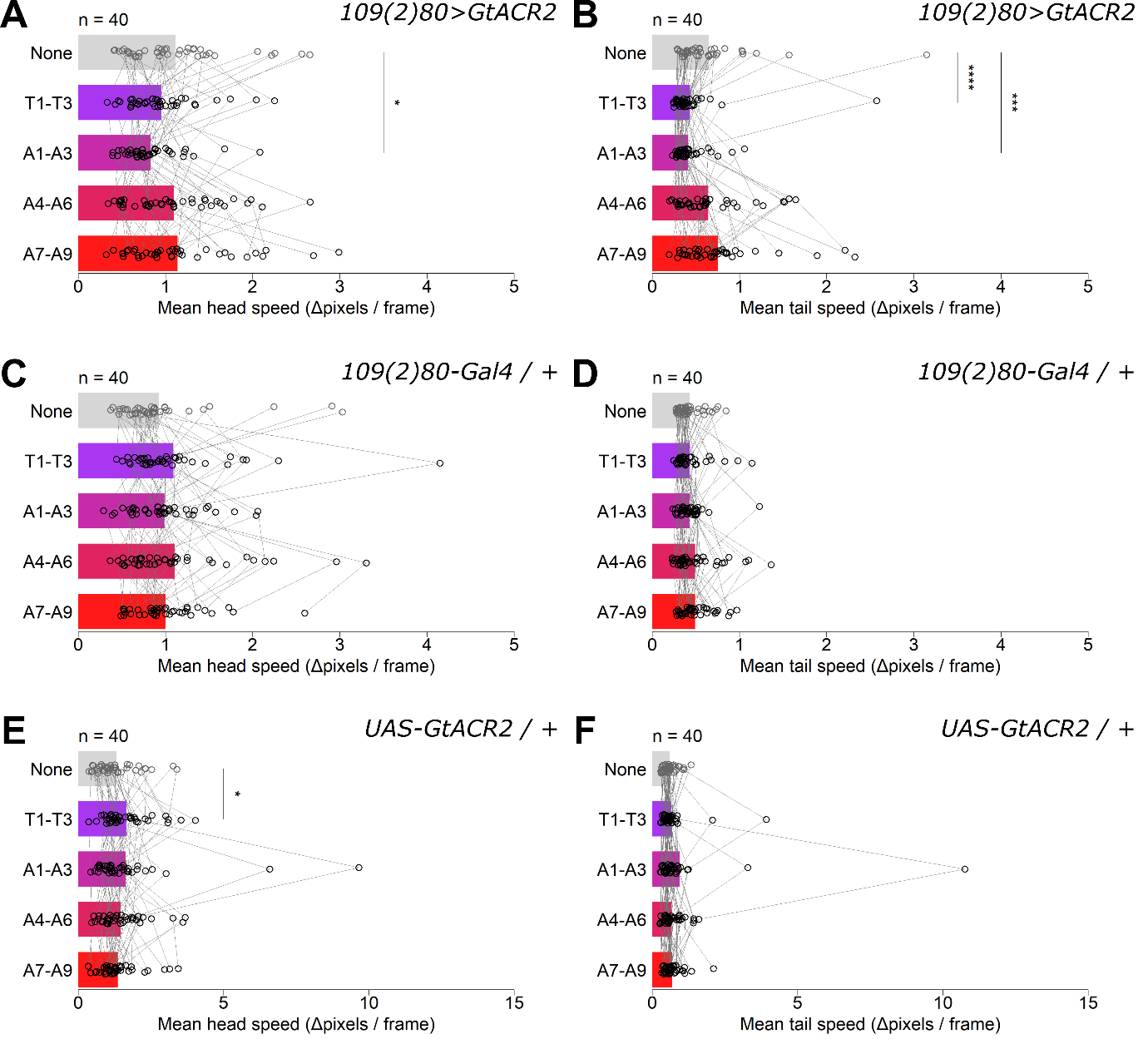

**Figure S4. Mean head and tail speeds under regional illumination. A)** Mean head speed for third instar larvae of the 109(2)80>GtACR2 genotype. **B)** Mean tail speed for third-instar larvae of the 109(2)80>GtACR2 genotype. **C)** Mean head speed for third instar larvae of the 109(2)80-Gal4 genotype. **D)** Mean tail speed for third-instar larvae of the 109(2)80-Gal4 genotype. **E)** Mean head speed for third instar larvae of the UAS-GtACR2 genotype. **F)** Mean tail speed for third-instar larvae of the UAS-GtACR2 genotype. In all plots, the bars show mean values, and points show individual measurements with measurements from the same larva being joined by grey lines. Analysis of deviance for a mixed model predicting head speed from genotype and illumination condition revealed a significant interaction effect (F_(8, 468)_ = 2.07, P = .037, n = 40). Similarly, analysis of deviance for an analogous mixed model predicting tail speed also revealed a significant interaction effect (F_(8, 468)_ = 4.39, P < .001 n = 40). ‘*’ = P < .05, ‘***’ = P < .001 and ‘****’ = P < .0001 for post-hoc two-sided comparisons between conditions of light illumination, following Dunnet’s approach with Sidak’s adjustment for multiple comparisons.

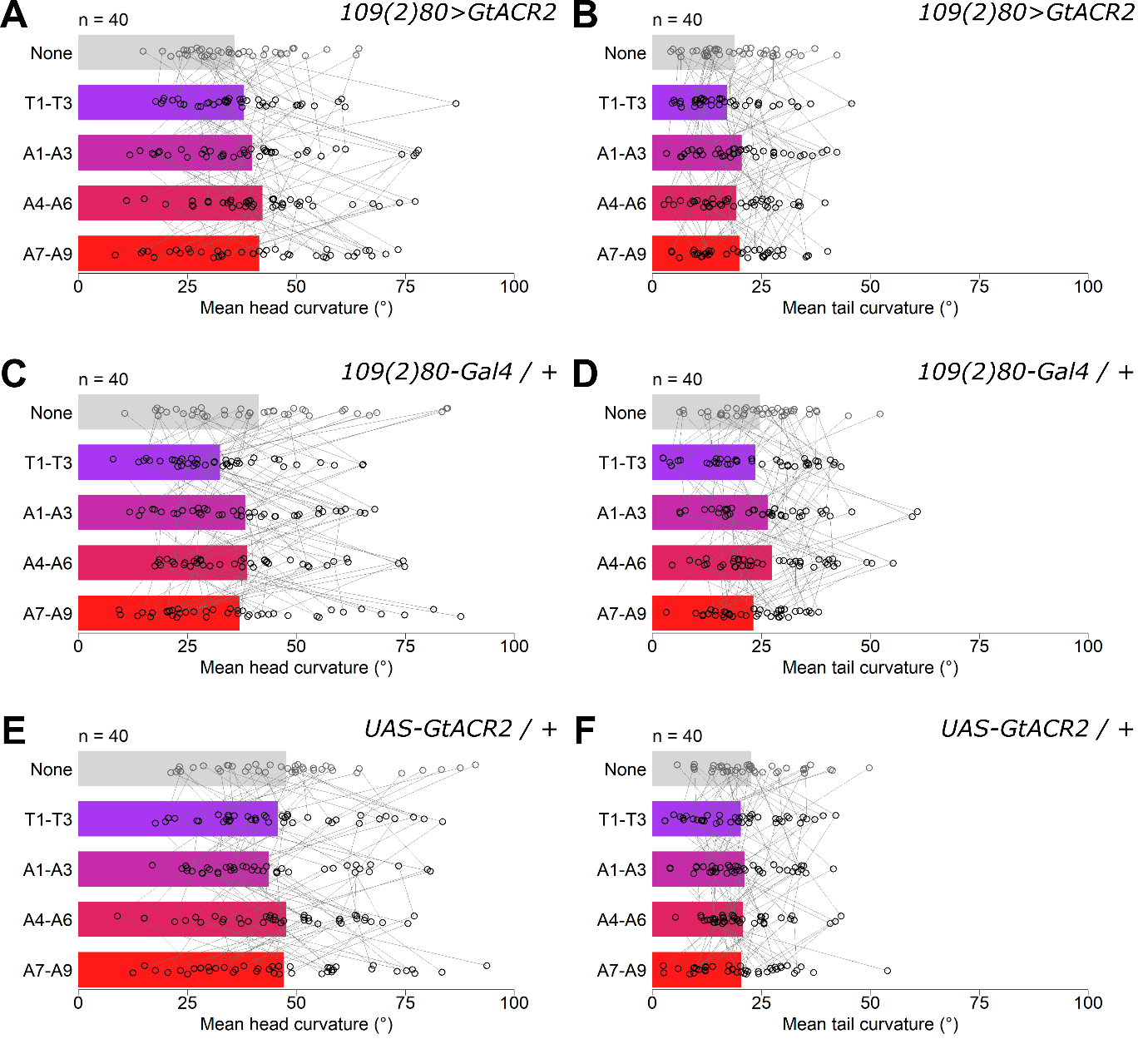

**Figure S5. Mean head and tail absolute curvature angles under regional illumination.** **A)** Mean head curvature for third instar larvae of the 109(2)80>GtACR2 genotype. **B)** Mean tail curvature for third-instar larvae of the 109(2)80>GtACR2 genotype. **C)** Mean head curvature for third instar larvae of the 109(2)80-Gal4 genotype. **D)** Mean tail curvature for third-instar larvae of the 109(2)80-Gal4 genotype. **E)** Mean head curvature for third instar larvae of the UAS-GtACR2 genotype. **F)** Mean tail curvature for third-instar larvae of the UAS-GtACR2 genotype. In all plots, the bars show mean values, and points show individual measurements with measurements from the same larva being joined by grey lines. Analysis of deviance for a mixed model predicting head speed from genotype and illumination condition found a non-significant interaction effect (F_(8, 468)_ = 1.06 , P = .392, n = 40). Similarly, analysis of deviance for an analogous mixed model predicting tail speed also found a non-significant interaction effect (F_(8, 468)_ = 0.68, P = 0.710, n = 40). Post-hoc two-sided comparisons between conditions of light illumination following Dunnet’s approach found no significant differences between the groups.

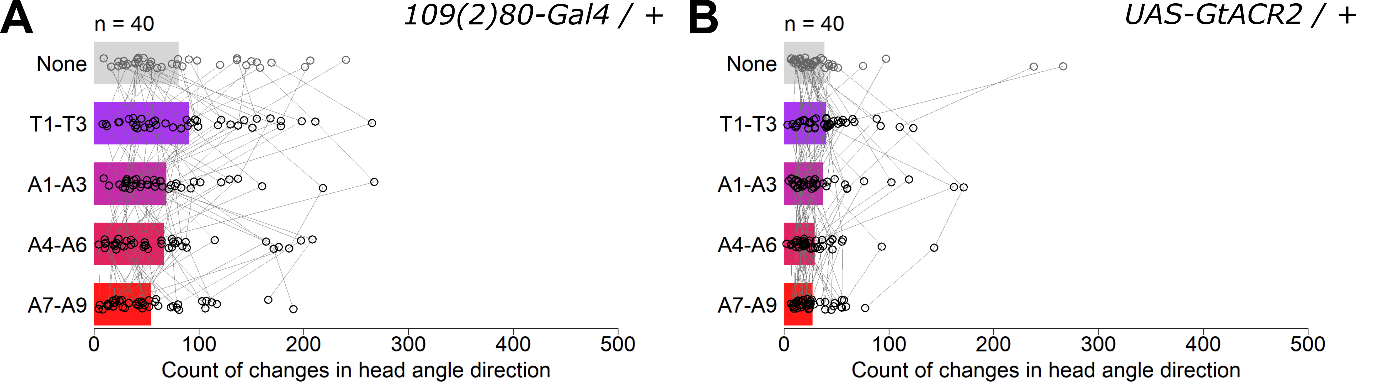

**Figure S6. Counts of changes in head angle direction under regional illumination for genetic control lines. A)** Counts of changes in head angle direction across the five illumination conditions for third instar larvae of the 109(2)80-Gal4 genotype. The bars show mean values, and points show individual measurements with measurements from the same larva being joined by grey lines. **B)** As A, but for larvae of the UAS-GtARC2 genotype. No statistical increases were observed between the control condition and the four illumination conditions.

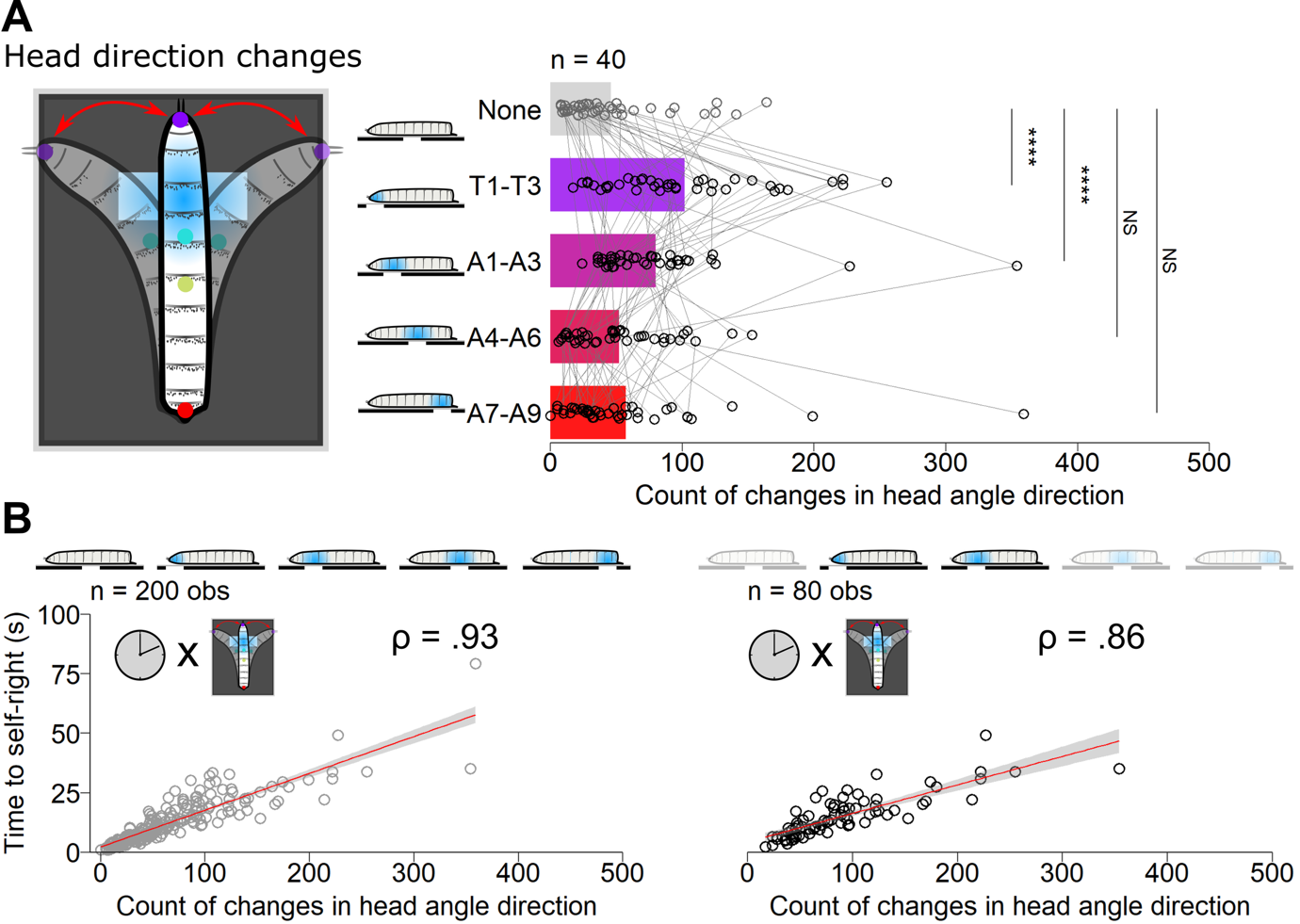

**Figure S7. Head casting patterns under localised optogenetic inhibition of daIV sensory neurons along the anterior-posterior axis.** A) Counts of the changes in head curvature direction under localised optogenetic inhibition of daIV neurons. The counts were calculated as the number of times a larva went from bending towards one direction with the head to bending in the other direction. Bars show the mean values for each condition, while points show counts for individual samples with measurements from the same larva being connected by lines. Analysis of deviance for a negative binomial model including control genotypes revealed a significant interaction of illumination condition and genotype (Χ2(8) = 30.61, P < .001, n = 40). ‘****’ = P < .0001 for post-hoc one-sided comparisons between conditions of light illumination, following Dunnet’s approach with Sidak’s adjustment for multiple comparisons. B) Correlations between count of head direction changes and self-righting times, for all illumination conditions (left) and just the anterior illumination conditions (right) in ppk>GtACR2 larvae. Points show individual observations, while the red line indicates a linear regression. P < .0001 for both Spearman correlations. Note that in all panels, one observation is not shown to improve visualisation (value = 1063 counts).

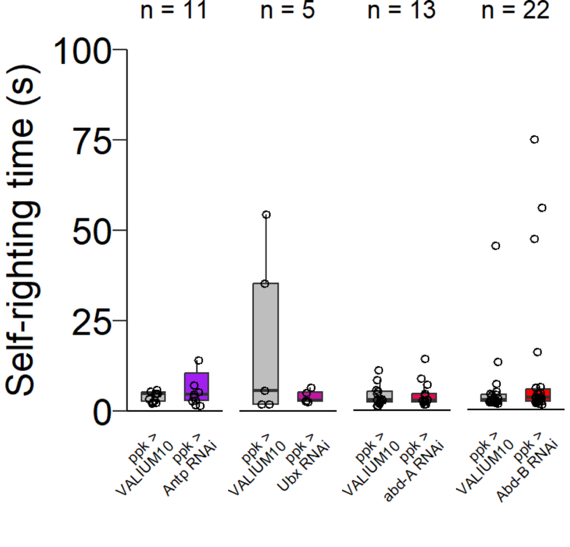

**Figure S8. Reduction of Hox gene expression within the ppk domain does not affect SR time.** Self-righting times of first instar larvae expressing a Hox gene RNAi construct in the ppk domain.

LEGENDS FOR MOVIES

**Video S1: The opto-axial technique for regional optogenetics.** The video shows a bird’s-eye perspective of the full opto-axial technique for regional sensory inhibition during self-righting. The larva has been positioned on a dry coverslip such that segments A1-A3 align with the arena slit. The blue LED under the arena is activated, and then the larva is freed to move by delivery of moisture using a fine paintbrush. The recording continues until after the larva has completed self-righting, at which point the tracheal trunks are visible along the dorsal side.

**Video S2: Postural tracking and feature extraction from the opto-axial technique.**

The video shows the same experiment as Video S1, following analysis using DeepLabCut. The video has been trimmed so the start coincides with the moisture delivery (water unlocking) and the end coincides with the precise completion of self-righting. Points on the larva indicate the maximum-likelihood positions of the head (purple), anterior middle (cyan), posterior middle (yellow) and tail (red) as estimated by DeepLabCut. The text labels indicate in degrees the calculated curvature angles of the head and tail for each frame of the video (see methods for details).
